# Mild perinatal hypoxia uncouples excitatory–inhibitory circuit maturation and reprograms neocortical organization

**DOI:** 10.64898/2026.05.07.723460

**Authors:** Matea Drlje-Ćurt, Sara Trnski-Levak, Siniša Škokić, Davide di Censo, Mihaela Bobić-Rasonja, Eugene Kim, Ivona Kirchbaum, Andrija Štajduhar, Katarina Ilić, Diana Cash, Miloš Judaš, Nataša Jovanov-Milošević

## Abstract

Perinatal hypoxia is a major contributor to neurodevelopmental disorders; however, the consequences of mild-to-moderate perinatal hypoxia (MPH) remain insufficiently characterized. Here, we investigated cortical plasticity following MPH using a multimodal approach that combines behavioral assessment, histological analysis, and in vivo magnetic resonance imaging (MRI).

Fifty-six Wistar Han rats were exposed to hypoxia or normoxia at postnatal day 1 (P1). Neurodevelopmental assessment from P3 to P14 revealed impaired rooting and vibrissae-placing reflexes in hypoxic rats. Histological analysis demonstrated: altered expression of microtubule-associated protein-2, apical dendrite bundling, reduced neurofilament-H expression, and decreased dendritic arbor complexity in large pyramidal neurons, indicating disrupted maturation of excitatory circuits. Increased parvalbumin expression, higher interneuron density, and its enhanced neurite elaboration indicated precocious development of inhibitory circuits, consistent with a compensatory response.

MRI at P15, combined with whole-brain voxel-wise analysis, revealed a significant increase in fractional anisotropy in the anterior cingulate cortex (ACC). Convergent behavioral, histological, and imaging findings identified the ACC as the most vulnerable region following MPH, followed by the somatosensory cortex.

These findings reveal early cytoarchitectural and MRI detectable correlates of a single episode of MPH, which, together with previous findings from this model, support the neurodevelopmental origin of persistent alterations in cortical structure and circuit function, characterized by an excitatory-inhibitory imbalance. The study identifies and defines a framework for understanding region-specific vulnerability and plasticity in the immature brain, with implications for improving the early detection of subtle perinatal brain injury, as a prerequisite for timely therapeutic intervention.

## INTRODUCTION

Premature birth complicated by perinatal hypoxia and/or ischemia (HI) remains a major cause of neonatal brain injury (Volpe 2008; Volpe 2009; Beacom et al. 2025; Chakkarapani et al. 2025). Mild-to-moderate perinatal hypoxia (MPH) represents a particularly challenging and under-recognized form of HI. Although infants affected by MPH may initially appear clinically unaffected, impairments often emerge later, with cognitive and behavioral abnormalities reported in 30–40% of preterm-born children at school age (Voss et al. 2007; Allen 2008; Saigal and Doyle 2008; Neubauer et al. 2008). During the critical developmental period in humans, from 23-32 gestational weeks, histogenetic processes in the brain are tightly regulated and include axonal elongation, dendritic arborization, cortical layer differentiation, and synaptogenesis, accompanied by synaptic pruning and specialization (Johnston Michael V. 1998; Kostović and Jovanov-Milošević 2006; Kostović and Judaš 2010; Mrzljak et al. 1990). Disruption of these processes by HI leads to region- and cell type-specific vulnerability, depending on developmental stage and injury severity (Johnston Michael V. 1998; McQuillen et al. 2003). In preterm-born individuals, long-term consequences include white matter abnormalities (Nosarti et al. 2002; Marlow et al. 2005), attributed to the vulnerability of oligodendrocyte progenitors to HI (Hüppi et al. 1996; Morken et al. 2013; Gopagondanahalli et al. 2016). Additionally, the vulnerability of subplate neurons (SpNs), which form one of the earliest functional cortical circuits and play a critical role in the establishment and maturation of thalamo-cortical connections, in both humans and rodents, is underscored (Kostović and Molliver 1974; Chun et al. 1987; Kostovic and Rakic 1990; Kostović and Jovanov-Milošević 2008; Friedlander 2009; Kanold 2009; Kanold and Luhmann 2010; Wang et al. 2010; Kostović 2020; Molnár and Kwan 2024).

Despite these challenges in perinatal medicine, studies of MPH remain limited in both humans and animal models (Clowry et al. 2014; Millar et al. 2017), partly due to the long-standing assumption of a favorable neurological prognosis (Conway et al. 2018; Li et al. 2022). As a result, early injury may remain unrecognized or be detected late, reducing the opportunity for timely therapeutic intervention. Furthermore, early structural correlates of MPH, particularly those detectable by translational imaging methods such as magnetic resonance imaging (MRI), remain insufficiently characterized, especially in rodent models. Regardless of species differences, key milestones of cortical differentiation and maturation are evolutionarily conserved and broadly comparable between humans and rodents (Workman et al. 2013; Semple et al. 2013). Rodent models are therefore widely used to investigate mechanisms underlying neurodevelopmental disorders (Molnár et al. 1998; Van De Looij et al. 2011; Semple et al. 2013; Jisa et al. 2018; Luhmann and Fukuda 2020; Luhmann 2023). The most commonly used rodent model of HI involves induction of severe injury at postnatal day 7 (P7), as well as modifications at earlier stages (e.g., P2–P3) (Rice et al. 1980; Vannucci et al. 1999; Sizonenko et al. 2003; Vannucci and Back 2022). However, models specifically addressing mild forms, such as MPH, remain under-researched.

MRI provides a non-invasive approach for assessing structural brain alterations in both clinical and experimental settings (Van De Looij et al. 2014). In humans, MRI studies have demonstrated reduced cerebral volumes and region-specific vulnerability along a rostro-caudal developmental gradient (Thompson et al. 2007; Bregant et al. 2013). In contrast, comparable patterns of regional vulnerability following MPH have not been established in rodent models. To address this, we applied a multimodal approach combining early neurodevelopmental behavioral testing, histological analysis of cortical organization, and in vivo MRI following MPH, to examine early cortical vulnerability and plasticity. We previously showed that this model exhibits long-lasting abnormalities that persist in the mature brains of adult rats (Trnski et al. 2022; Nikolic et al. 2023). We therefore hypothesized that hypoxia induced at P1 in rats, which broadly corresponds to a 23–32 gestation week in humans, more closely reflects mild hypoxic injury in premature infants with respect to cortical developmental processes (Mallard and Vexler 2015). We aim to identify structural and MRI-associated correlates, or imaging biomarkers, of MPH in a rat model, providing a research tool to improve prognostic and therapeutic assessment in human perinatal brain injury.

## MATERIALS AND METHODS

### Animals and housing

All animal experiments complied with the ARRIVE guidelines and have been carried out following the United Kingdom Animals (Scientific Procedures) Act 1986. Maximum effort was made to reduce the number of animals used and minimize animal discomfort. In total, fifty-six 1-day-old Wistar Han (RccHan: WIST) rats were obtained from our breeding facility (School of Medicine, University of Zagreb, Croatia). Rats of an average body mass (6.83 ±0.68g) were randomly assigned to either the hypoxic or control group, maintaining equal sex representation. The day of birth was considered P0 until noon of the next day when P1 began. After exposure to hypoxia, pups from both groups were permanently marked by a toe tattoo (NEO-9 Neonate Tattoo System, AgnTho’s AB, Sweden). Immediately after the hypoxic exposure twelve rats (6 hypoxic and 6 control) were sacrificed to determine the Acid-Base Status (ABS). For neurodevelopmental behavioral and reflex testing in postnatal rats, twenty-four (12 hypoxic and 12 control) rats were used, and the testing was performed from P3 to P14. Three males from the control group died during the testing period, and their testing results were excluded from statistical analysis. At P15, a subset of rats (8 hypoxic and 8 control) was scanned using MRI and later sacrificed for histological and immunohistochemical analysis. Additional rats were used, comprising a larger cohort of forty-four P15 rats (22 hypoxic and 22 control), for histological and immunohistochemical analyses. Supplementary Table 1 displays a list of experiments and the total number of rats used in the study. The study design is presented in Supplementary Fig. S1.

Rats were housed in polysulfone cages within a controlled environment with a temperature of 21 ± 2°C and humidity maintained at 65 ± 5%. A 12h:12h light: dark cycle was employed with the light period beginning at 7 am. Food (4RF21C, Mucedola srl, Settimo Milanese MI, Italy) and tap water were provided ad libitum.

### Hypoxia-inducing protocol

The P1 pups were weighed, sex determined and separated from dams. The hypoxic group (4 female (F)+4 male (M) pups per session) was placed in a closed-heated hypoxic chamber (STEMCELL Technologies Inc. Vancouver, Canada, Cat. No. 27310) connected to the gas cylinder containing the gas mixture of 8% O_2_ and 92% N_2_ (Messer Croatia Plin d.o.o., Zapresic, Croatia), using a Spectromed cylinder pressure regulator FM41-S1 (Spectron Gas Control Systems GmbH, Langen, Germany), which enabled a continuous flow of a mixture of gases of 3.5 L/min for 2 hours. The chamber was filled with bedding from cages where pups are housed with dams. The humidity, temperature, and oxygen level within the chamber were monitored using an optical oxygen gas sensor FDO_2_ (PyroScience GmbH, Aachen, Germany) connected to a computer. The CO_2_ adsorbents were placed in the chamber to absorb exhaled CO_2_. The chamber was placed on the heating pad to maintain a constant temperature of approximately 30°C. The control group (4F+4M per session) was exposed to room air (21%O_2_, 79%N_2_ for 2h), with other parameters the same as the hypoxic group. Rat pups were permanently tattooed and returned to their cages and dams. The blood ABS assessment in rat pups after this type of treatment was previously reported by our group (Trnski et al. 2022). The ABS measured in the peripheral blood of rats in this study immediately after the hypoxia treatment at P1 confirmed lactic acidosis (p=0.002, Mann-Whitney). A systemic physiological response with a sufficient compensatory capacity was evident by increased pH (p=0.007, Mann-Whitney) and decreased bicarbonate (p= 0.009, Mann-Whitney) values in the blood (Supplementary Fig. S2).

### Early neurodevelopmental behavior assessments

Neurodevelopmental behavioral and reflex testing was performed longitudinally on each rat from P3 to P14 daily between 11 am and 2 pm. Rats were housed with dams, and litter size on the day following birth was standardized to 8 pups per litter. Each day, rat pups were separated from dams for 3 hours and randomly selected for individual testing by two blinded researchers to the experimental group. All rats were returned to their dams in the home cage simultaneously after testing. For neurodevelopmental milestones, physical landmarks were observed: pinnae detachment, eye opening, incisor eruption, fur development, descending testis (in males), and body weight. During the 3-minute observation period, the development of quadruped stance–posture, head, forelimbs, and shoulders elevation, and quadruped locomotion – pivoting, crawling, walking was also observed. Ambulation, surface, and air righting tests were used for general motor function development. Negative geotaxis, cliff avoidance, and rooting reflex were used for sensory-motor development. Vibrissae placement and tactile startle reflexes were observed for somatosensory development. All tests were performed as described in the literature (Fox 1965; Smart and Dobbing 1971; Altman and Sudarshan 1975; Heyser 2003; Feather-Schussler and Ferguson 2016; Nguyen et al. 2017).

### MR Imaging

The in vivo structural and diffusion MRI protocol (Romanelli et al. 2022; Justić et al. 2022) was performed on a 7T Bruker BioSpec small animal scanner equipped with Paravision software support (BioSpec 70/20 USR with Paravision 6.0.1. software version, Bruker Biospin, Germany). The Tx/Rx configuration, using an 86 mm transmit volume coil (MT0381, Bruker Biospin, Germany) for transmitting (Tx) and a 2-element mouse brain surface coil (MT0042, Bruker Biospin, Germany) for receiving (Rx), was used for all recordings. The rats were anesthetized with 4% Isoflurane (Isofluran-Piramal, Piramal Critical Care, Germany) in a 30/70 oxygen-nitrogen mixture. The anesthetized animals are then placed in the appropriate MRI cradle. During imaging, body temperature and breathing frequency were continuously monitored and measured by the Medres MLT (Medres, Cologne, Germany) physiological system. Adjusting the amount of anesthetic in the mixture maintained stable anesthesia and a breathing rate at 45-50 breaths per minute. The scan protocol consisted of voxel-wise maps of T1 and T2 relaxation times (FOV 16/12 mm, 125 µm isotropic in-plane resolution producing an image size of 128 x 96 pixels, 21 slices with 700 µm slice thickness and 200 µm slice gap), 3D anatomical scans and a diffusion tensor imaging (DTI) (FOV 12/16/13.5 mm, 125 µm in-plane resolution producing an image size of 96 x 128 pixels, 27 slices with 500 µm slice thickness). The acquired anatomical scans were T1-weighted inversion recovery (T1-IR), T2-weighted, and two T2*-weighted scans. The diffusion scan was performed with a DTI-EPI sequence. Optimal field homogeneity for the DTI-EPI scan was achieved by additional localized shimming using the MAPSHIM algorithm (see Supplementary Table 2 for detailed scan parameters).

### Image processing and analysis

After visual inspection, one control subject was excluded due to strong distortion artifacts in the DTI images. All other subjects were processed using the following pipeline: denoising (dwidenoise), deringing (mrdegibbs) performed with Mrtrix3 (Tournier et al. 2019), and DTI fitting (dtifit) performed with FSL (Avants et al. 2011; Jenkinson et al. 2012). A multi-parametric study-specific DTI template was created with the fractional anisotropy (FA) and mean diffusivity (MD) maps using the antsMultiVariateTemplateConstruction2.sh script in ANTs. Sequential rigid-body, affine, and SyN registrations were performed using antsRegistration to align all subjects to the study-specific template, which was then aligned to the P14 rat brain atlas created by Rumple et al. (Rumple et al. 2013).

The IR-RARE, T2w-RARE, and MSME images were used to perform tensor-based morphometry to test for regional brain volume differences between the groups. As above, each subject was registered to a study-specific template created from these three image types. Voxel-wise volumes were quantified by calculating maps of the Jacobian determinants of the resultant deformation fields.

Voxel-based analysis was performed using permutation tests (Winkler et al. 2014) and threshold-free cluster enhancement (TFCE) to test for differences in diffusion metrics and log-transformed Jacobian determinants between the hypoxic and control groups. The results are presented using the dual-coding approach (Allen et al. 2012; Taylor et al. 2023): the effect size, i.e., the magnitude of the difference between groups, is represented by the overlay color. The overlay opacity increases with statistical significance, i.e., lower TFCE-corrected p-values. The p-values were derived from the statistical parametric maps. Additionally, black contours denote clusters that exceed the significance threshold p < 0.01.

### Histological and immunohistochemical methods

For immunohistochemistry, the P15 rats were deeply anesthetized with an intraperitoneal administration mixture of 80 mg/kg ketamine and 10 mg/kg xylazine (both from Bioveta, Czech Republic) in an isotonic solution (Ramsey 2017). After confirming that the rats are under deep anesthesia (loss of medial and lateral palpebral reflexes and loss of deep pain sensation tested by pressing forceps on the posterior third of the tail and pressing the skin between the fingers), rats were transcardially perfused (Gage et al. 2012) with ice-cold phosphate-buffered saline (PBS), subsequently with 4% paraformaldehyde (PFA in 0.1 M PBS; pH = 7.4) and then decapitated. Following 24-hour fixation in 4% PFA in 0.1 M PBS, brain samples were cryoprotected by being placed in increasing sucrose concentrations (20%, 30%) and then snap-frozen in isopentane cooled to -80°C. The brains were further embedded in paraffin and sectioned in the coronal plane to a thickness of 10 μm. Deparaffinization was performed using a descending series of alcohols (xylene, 100% ethanol, 96% ethanol, 70% ethanol), and the sections were then washed in distilled water. Adjacent sections were processed by Cresyl-violet (Nissl) and Mowry staining to delineate histoarchitectural boundaries and cortical layers of anterior cingulate (ACC) and somatosensory cortex (Paxinos and Watson 2014; Vogt and Paxinos 2014; Pallomero-Gallagher and Zilles 2015; Swanson 2018). A 0.5% Cresyl violet stain (Chemika, Girraween, Australia) in distilled water at a 1:2 ratio was used for 3 minutes. The sections were then washed in distilled water and dehydrated through an ascending series of alcohols (70% ethanol with a few drops of 10% acetic acid (CH₃COOH), 96% ethanol, 100% ethanol) and finally cleared with xylene and then Histo-Clear solution (Chemika, Girraween, Australia). Changes in hyaluronic acid, an extracellular matrix (ECM) component, were evaluated using the Mowry method (Luna 1960; Jovanov-Milošević et al. 2014; Bobić Rasonja et al. 2019). After deparaffinization and rinsing in 12% acetic acid, the sections were left for 1h in a working solution of colloidal iron (an aqueous solution of colloidal iron in concentrated acetic acid), then rinsed four times in 12% acetic acid. The next step was incubation for 20 minutes in a solution of 5% potassium ferrocyanide and 5% hydrochloric acid, followed by rinsing in running and distilled water. After immersion in 96% and 100% ethanol and BioClear (BioGnost) the sections were coverslipped with BioMount (BioGnost) mounting medium.

Adjacent coronal sections were processed immunohistochemically as follows: after dewaxing in xylol and rehydration in a graded series of ethanol solutions, sections were washed in PBS, antigen retrieval in heated citrate buffer (pH=6.0) was performed, sections were cooled for half an hour at room temperature (RT) and were rinsed in PBS. Endogenous peroxidase activity was blocked by 0.3% H_2_O_2_ in a 75 % methanol solution. Sections were rinsed in PBS and immersed for 1h in the blocking solution (PBS containing 5% bovine serum albumin and 0.5% TritonX-100), all from Sigma (St. Louis, MO) at RT to intercept non-specific staining. Incubation with primary antibodies was performed for 48 hours at 4°C for all antibodies: Anti-Microtubule-associated protein 2 (MAP2) (Sigma),Anti-Neurofilament-H(NF-H) (Biolegend), Anti-Parvalbumin (PV) (Abcam), Anti-Myelin Basic Protein (MBP), Anti-Complexin 3 (Cplx3) (Synaptic Systems), Anti-Keratan Sulfate (KS), (MEMD Milipore Corp) Anti-*Wisteria floribunda* agglutinin (WFA)(Sigma-Aldrich), Anti-Lumican (Abcam), Anti-Neurocan (Sigma-Aldrich), Anti Versican (Thermofisher) (see Supplementary Table 3 for details). Following incubation with an appropriate secondary antibody for 2h at RT (Vectastain ABC kit, Vector Laboratories, Burlingame, USA), sections were further incubated in Vectastain ABC reagent (streptavidin-peroxidase complex) for 1h at RT and rinsed in PBS. Peroxidase activity was visualized with Ni-3,3-diaminobenzidine (Sigma D0426, St. Louis, MO, USA). After drying at RT, sections were cleared in BioClear (BioGnost) and coverslipped with BioMount (BioGnost). Negative controls were included in all immunohistochemical experiments by replacing the primary antibody with the blocking solution.

### Histological and immunohistochemical analyses

Qualitative analysis of the stained histological sections was performed using an Olympus BX53 light microscope, and images were captured with an Olympus UC-90 digital camera (Olympus Corporation, Shinjuku, Tokyo, Japan) or using a high-resolution digital slide scanner Nano-Zoomer 2.0RS (Hamamatsu, Japan). The quantifications of MAP2 and Mowry staining intensity, and the counting of PV and Cplx3-positive neurons were performed in ACC or somatosensory cortex at the approximate bregma levels 2.28 and 1.28. mm, respectively, (Paxinos and Watson 2014). For the quantification of the diffuse MAP2 staining (20 rats per group) of cortical layer I (the strongest immunoreactivity) and corpus callosum (minimum immunoreactivity), the region of interest (ROI) was delineated using Fiji software (FIJI, version Java 1.8.0_172 (64-bit). In addition, the mean staining intensity was measured, with pixel values ranging from 0 (maximum immunoreactive staining) to 255 (no immunoreactive staining) for each ROI. The mean gray value of ROI intensity for each section was calculated and normalized to the mean gray value intensity of the corpus callosum (minimum immunoreactivity). Afterward, the calculated ratios were divided by the maximum value to obtain an inverted scale ranging from 0 (minimum) to 1 (maximum). The quantitative analysis of the number of PV-positive interneurons (N=12 rats per group) was performed using Neurolucida 10 (MBF–Bioscience, Williston, ND, United States), and an Olympus BX61 microscope as described previously (Bicanic et al. 2017; Trnski et al. 2022). Semi-automated quantification of the number (16 rats per group) and intensity of immunoreactivity in Cplx3-positive neurons was performed using a custom-made macro in Fiji (ImageJ; Java 1.8.0_172, 64-bit). Contrast Limited Adaptive Histogram Equalization (CLAHE) was applied to all images (block size = 127, histogram bins = 256, maximum slope = 3, fast mode) to enhance signal and facilitate thresholding. A fixed ROI was defined using identical parameters across all images. Neurons were subsequently manually identified, labeled, and counted using the Magic Wand tool in Fiji software (ImageJ; Java 1.8.0_172, 64-bit). The intensity of somatic staining was assessed in 100 randomly selected neurons, which were manually delineated, and the mean gray value was calculated. The quantification of the diffuse Mowry staining (18 rats per group) across cortical layers I–VI, analysis was performed in three ROIs: layer I, layers II–III, and layers V–VI using Fiji software (FIJI, version Java 1.8.0_172 (64-bit). In addition, the mean staining intensity was measured, with pixel values ranging from 0 (maximum immunoreactive staining) to 255 (no immunoreactive staining) for each ROI. All statistical tests were conducted using Prism (GraphPad Software, version 10.0. Inc., La Jolla, CA, United States). Normality was assessed using the Shapiro–Wilk test. For normally distributed data, Welch’s t-test was used for two-group comparisons. For non-normally distributed data, the Mann–Whitney U test was applied. Data are reported as mean ± SD or median (interquartile range), as appropriate. All statistical tests were two-sided, and a significance threshold of α = 0.05 was used. P-values were adjusted for multiple testing across layers using the Holm-Bonferroni method to control the family-wise error rate.

## RESULTS

### Behavioral testing reveals altered maturation of basic reflexes in rats following MPH

Observation of the rats after the hypoxia treatment revealed no differences in locomotion between groups. In contrast, the vibrissae placing response was significantly less pronounced in hypoxic rats than in controls at P3 (p=0.037) and P5 (p=0.003) (Fig. 1, b). At P9, there was a peak in vibrissae-placing performance in both groups. This reflex persisted until P14, when, in hypoxic rats, an indicative drop in performance (p=0.052) was observed compared to control rats. The rooting reflex was also significantly less pronounced in hypoxic rats than in controls at P5 (p=0.046), P6 (p=0.031), and P7 (p=0.036) (Fig. 1, c). At P7, there was a peak in rooting reflex performance in controls, whereas in the hypoxic group, it occurred at P8. Afterward, this reflex is gradually lost. At P5, hypoxic pups showed significantly higher surface righting scores than controls (p = 0.039; Fig. 1e). At P6, scoring adjusted for time was also significantly greater in the hypoxic group (p = 0.027; Fig. 1, a), indicating faster righting performance. Briefly, locomotor development was unaffected, but alterations in basic reflexes were present in hypoxic group.

**Fig. 1.**
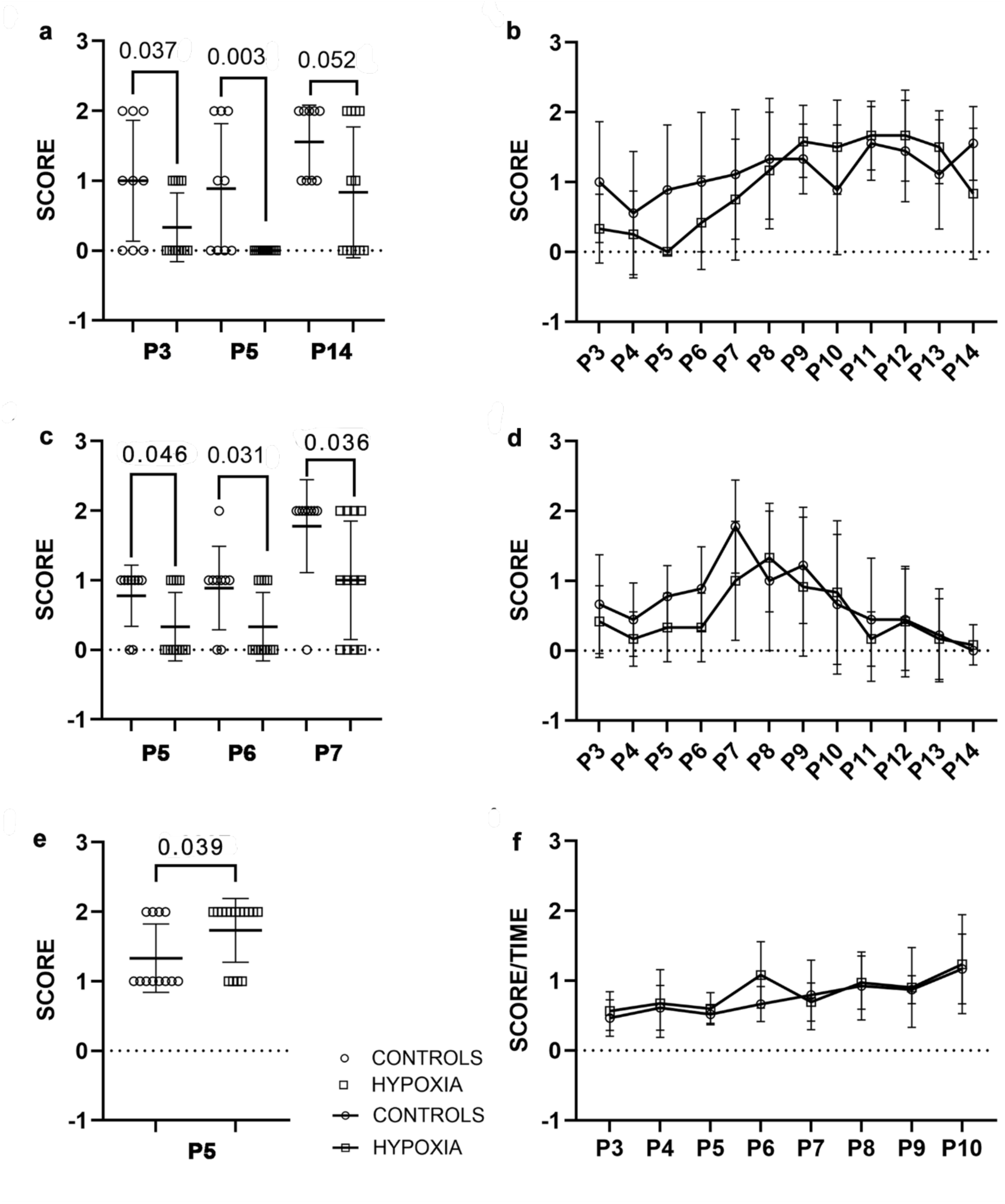
Sensory-motor and somatosensory development in rats from P3 to P14. Graphs display the results of the behavioral tests (a–f). Vibrissae placing response (a, b) was reduced in hypoxic rats at P3 (p = 0.037) and P5 (p = 0.003). Performance peaked at P9 in both groups and was maintained until P14, with a decline in hypoxic rats at P14 (p = 0.052). Rooting reflex (c, d) was reduced in hypoxic rats at P5 (p = 0.046), P6 (p = 0.031), and P7 (p = 0.036); the peak occurred at P7 in controls and P8 in hypoxic rats, followed by a gradual loss. In the surface righting test (e, f), hypoxic rats showed higher scores at P5 (p = 0.038) and P6 (p = 0.027), indicating faster performance. Data are shown as mean ± SD

### In vivo MRI detects the cortical lesions following MPH

T2-weighted imaging did not reveal any gross structural lesions or abnormalities in any of the examined rats. On the other hand, DTI imaging revealed clusters of increased FA in the anterior cingulate cortex (ACC; Fig. 2a), accompanied by decreased RD (Fig. 2b) and increased AD (Fig. 2c) in the corresponding brain regions of hypoxic rats compared with controls. The most significant clusters were at the Bregma level 1.28 mm (Paxinos and Watson 2014). For FA values of each rat, see Supplementary Table 4.

**Fig. 2.**
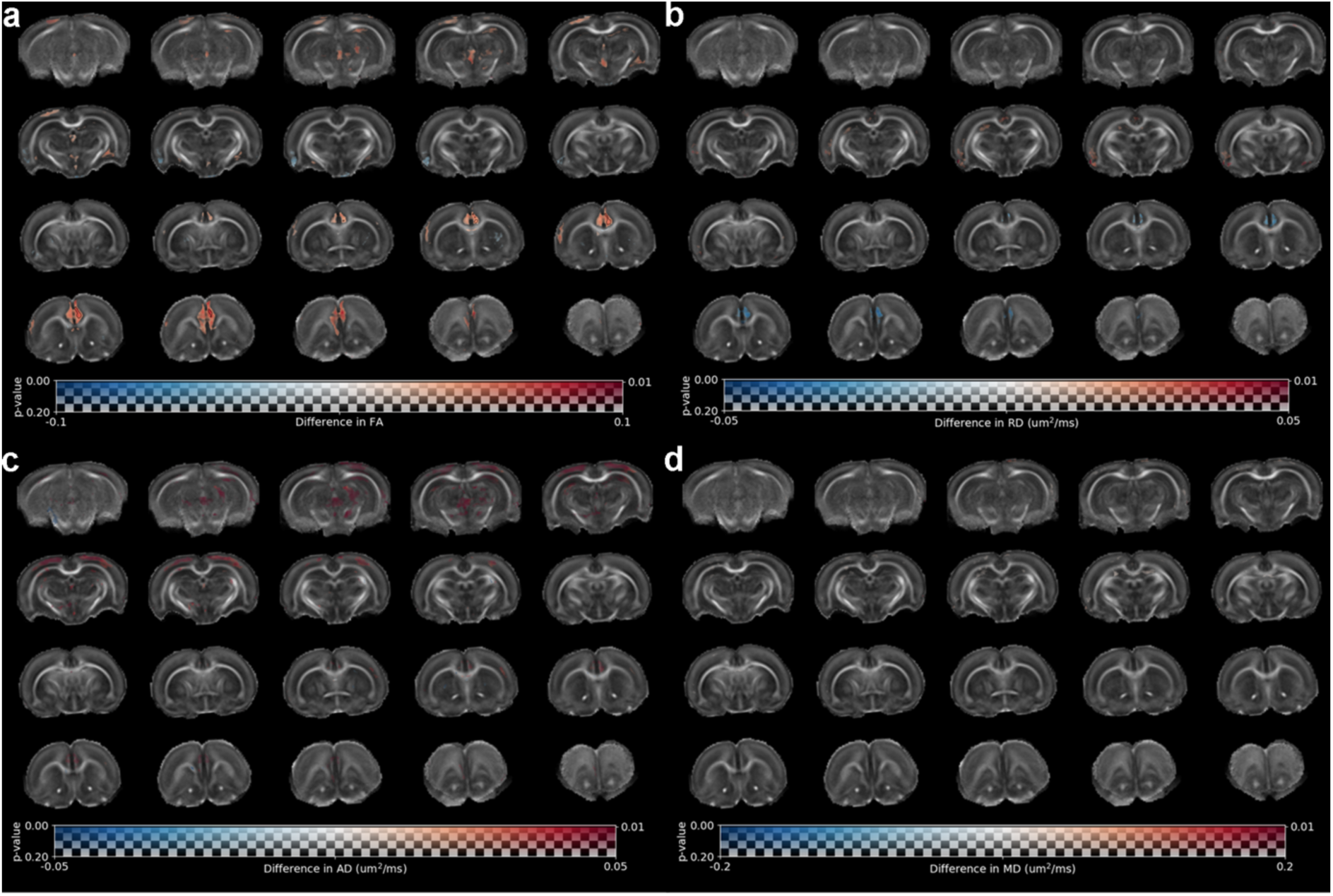
Whole-brain voxel-wise DTI differences between control and hypoxic rats at P15. Group differences (hypoxic vs controls) in fractional anisotropy (FA), radial diffusivity (RD), axial diffusivity (AD), and mean diffusivity (MD) (a–d) are shown as color overlays on a study-specific P15 rat DTI template. Red color indicates higher values in hypoxic rats; overlay opacity reflects statistical significance (greater opacity, lower p value). Black contours denote clusters with p < 0.01. Hypoxic rats showed increased FA in the prelimbic cortex and anterior cingulate cortex (ACC), with significant increase in the ACC (a). RD was reduced in the ACC, with small bilateral increases in the retrosplenial cortex (RSC) (b). AD was increased bilaterally in the ACC, motor, somatosensory, and visual cortices (c). No group differences were observed in MD (d)

The anatomical position of these changes corresponds to the precingulate region, anterior callosal radiations, cingulate bundle, and the deep (V, VI) layers of the ACC. An indicative increase in AD values (Fig. 2, c) and a trend toward increased FA (Fig. 2, a) in the supragranular layers of posterior dorsolateral cortex at Bregma levels around 0.48 (Paxinos and Watson 2014) were also detected.

Regional brain volume differences (Fig. 3) were found to be significant in the cortex (external) of the inferior colliculi (Fig. 3, red, circled in black). Volume differences were also observed in the dorsal thalamic nuclei and the sensory cortex, although they did not reach statistical significance (Fig. 3, red and orange). These regions showed a trend toward enlargement in hypoxic rats but did not differ in FA or other diffusivity parameters. Concomitantly, a bilaterally symmetrical decrease in the volume of reticular formation and midbrain pathways, although not statistically significant, confirms the affected sensory-motor-reflex circuit in hypoxic rats (Fig. 3, blue-green color scale).

**Fig. 3.**
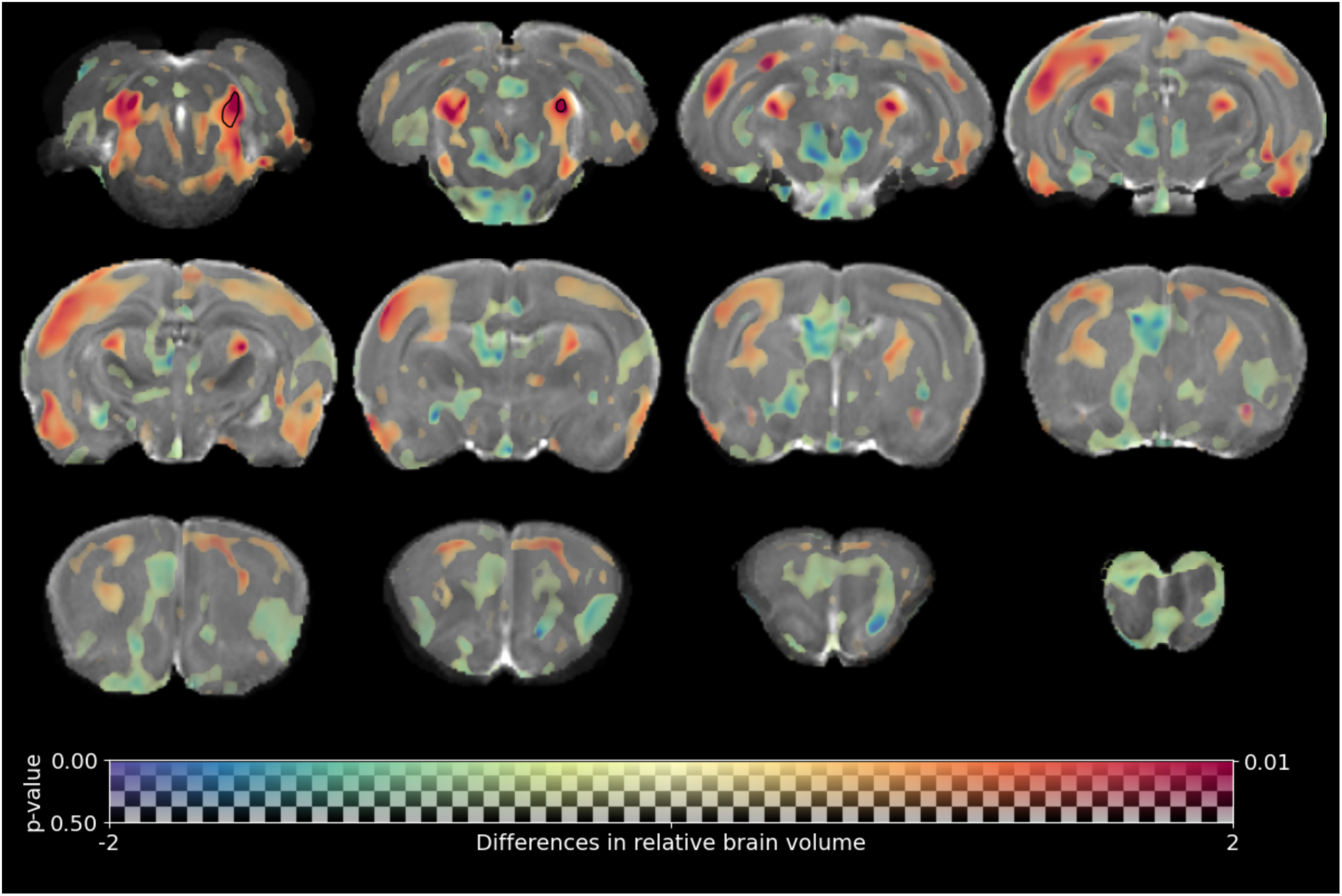
Whole-brain voxel-wise volumetric differences between hypoxic and control rats at P15. Voxel-wise differences in the volume determinant are shown as color overlays on a T2-weighted rat brain template (regions in red show increased volume in hypoxic rats compared to controls). Overlay opacity reflects statistical significance. Black contours denote clusters with p < 0.01. Hypoxic rats showed a trend toward increased cortical volumes, most prominently in the primary somatosensory, visual (primary and secondary), auditory (primary and secondary), and piriform cortices. Subcortical volume increases were observed in thalamic nuclei, including the lateral posterior and dorsal lateral geniculate nuclei, as well as in the external cortex of the inferior colliculus. Volume decreases were detected in the mesencephalon and pons, including the reticular formation, red nucleus, and raphe nuclei

### Neocortex structural aberrations post MPH

The qualitative analysis of whole brains and histological sections (Fig. 4-5, Supplementary Fig. S3), stained with the Nissl method, revealed preserved gross morphology, undisrupted cortical lamination and amount of ECM, no changes in the volume of major axonal tracts (anterior commissure, corpus callosum, internal capsule, optic tract), (Supplementary Fig. S5), no alteration of the volume or shape of the ventricles, and an absence of focal hypoxia-induced pathological features.

**Fig. 4.**
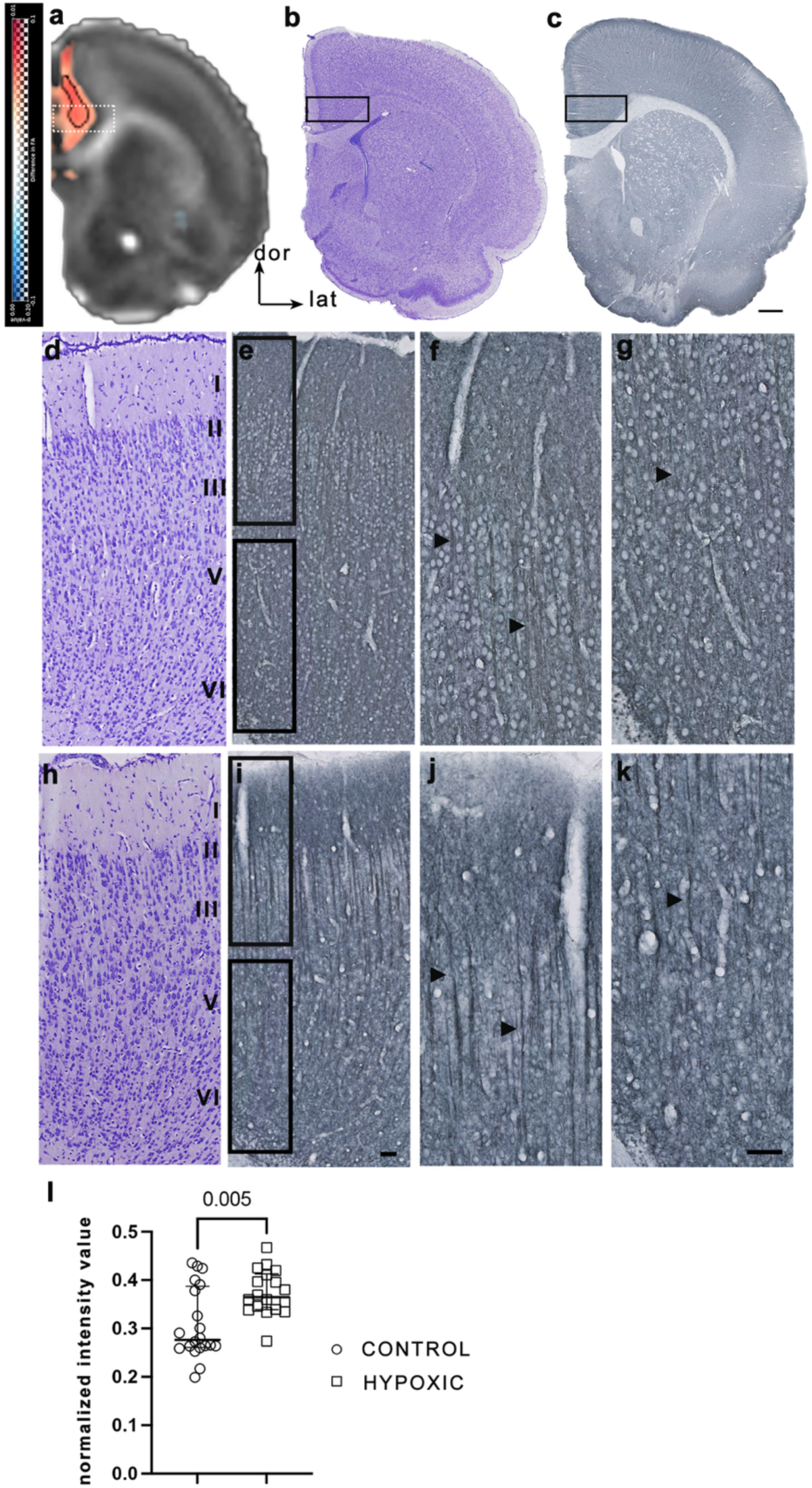
Changes in FA and MAP2 immunoreactivity in the ACC at P15 following MPH. Coronal sections at the level of the ACC (∼+2.28 mm from Bregma) show (a) group differences in FA and representative staining with (b) Nissl and (c) MAP2 immunohistochemistry. Black boxes in (b, c) indicate regions enlarged in (d–g) (control) and (h–k) (hypoxic). Nissl staining (d, h) shows preserved cytoarchitecture, cortical lamination, and indirectly the ECM amount in hypoxic rats, without gross pathological changes. MAP2 immunostaining (e–g, i–k) revealed altered immunoreactivity in the ACC. Staining in brains from the hypoxic group showed prominent, vertically oriented bundles of apical dendrites (i), compared to weaker staining and less distinct dendrites in controls (e). These differences are most evident in supragranular layers II–III (f, j; arrowheads) and are also present in infragranular layer V (g, k). Quantification (l) shows increased MAP2 immunoreactivity in layer I of the ACC (mean intensity normalized to the corpus callosum), presented as median ± IQR, with a significant increase in hypoxic rats compared to controls (p = 0.005, Mann–Whitney test). Scale bars: 0.5 mm in (c) applies to (b); 50 μm in (i, k) applies to (d–k)

**Fig. 5.**
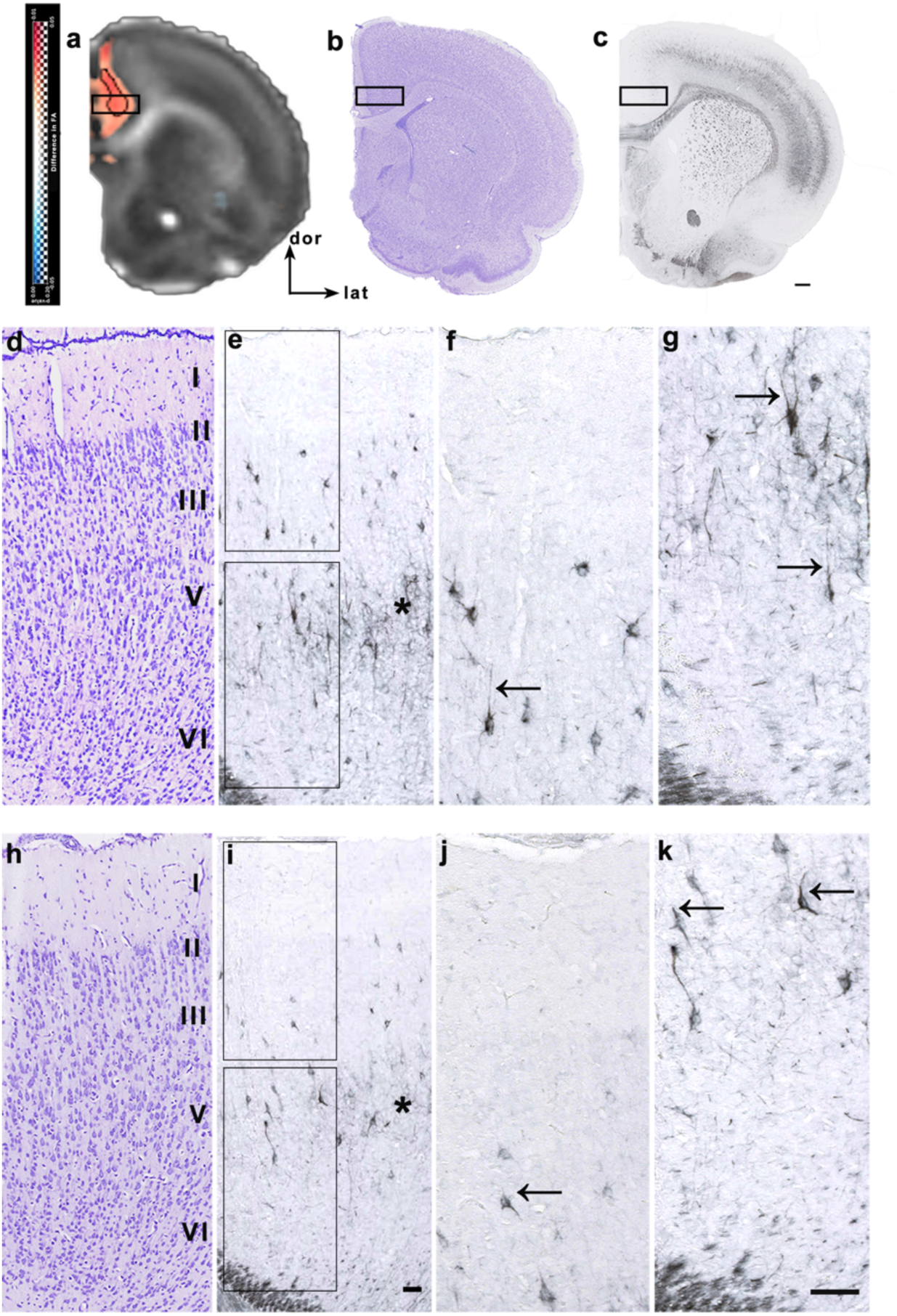
Changes in FA and NF-H expression in the ACC at P15 following MPH. Coronal sections at the level of the ACC (∼+2.28 mm from Bregma) show (a) group differences in FA and representative staining with (b) Nissl and (c) - NF-H. Black boxes in (b, c) indicate regions enlarged in (d–g) (control) and (h–k) (hypoxic). Cortical lamination and layer delineation were assessed using Nissl staining (d, h). Black rectangles in (e, i) indicate regions enlarged in (f, j) (supragranular layers) and (g, k) (infragranular layers). NF-H immunostaining revealed differences in the appearance of apical dendrites of excitatory projection neurons in layers III and V of the ACC in hypoxic rats (i–k) compared to controls (e–g). NF-H immunoreactivity was reduced in the layer V neuropil in hypoxic rats (asterisk in i) compared to controls (asterisk in e). In control rats, strong NF-H staining is present in apical and basal dendrites, indicating a more mature neuronal appearance. In hypoxic rats, only the proximal segment of the apical dendrite was strongly immunolabeled (arrows in j, k), compared to clearly stained and elongated apical dendrites in controls (arrows in f, g). Scale bars: 0.5 mm in (c) applies to (b); 50 μm in (i, k) applies to (d–k)

The MAP2 immunohistochemical staining, specific for neurons, revealed no differences or alterations in soma across cortical layers between control and hypoxic rats (Fig. 4). However, in hypoxic rats, significantly increased MAP2-positive fibers (presumably apical dendritic tufts) were observed in layer I of ACC. Additionally, the stronger MAP2 immunostaining of the apical dendrites, assembled in bundles, in the ACC cortex of hypoxic rats, revealed a more compact dendrite organization accompanied by reduced complexity of the dendritic tree (Fig.4). More specifically, MAP2 immunostaining revealed more tightly packed vertically oriented bundles of apical dendrites in hypoxic rats (Fig. 4, i-k), which appeared prominently MAP2-stained compared to the more dispersed staining in controls. A faint, dispersed, and more elaborate dendritic architecture was revealed in control rats (Fig.4, e-g). The differences in organization and stronger bundling of the apical dendrites in hypoxic rats were easily distinguished in the upper cortical layers, as the apical dendrites elongated through (Fig. 4, arrowheads in j), and in the deeper cortical layers (Fig. 4, arrowhead in k). Thus, control rats exhibited clear, diffuse MAP2 immunoreactivity in the cortical layers, marking the elaborated neuropil, with less intense staining in layer I (Fig. 4, f-g). In contrast, the MAP2 immunoreactivity in hypoxic rats was more concentrated in the apical dendrites of neurons in layers II and V and their bundles, and its distal dendritic tufts in layer I of ACC (Fig. 4, j-k). Quantitative analysis of MAP2 expression in layer I of the ACC corroborated the qualitative observations, revealing significantly higher MAP2 staining intensity in hypoxic rats (Fig. 4l).

The neurofilament-H specific (NF-H) immunohistochemical staining and its qualitative analysis revealed a less intense cellular and neuropil NF-H staining in the cortical layers III to V in the brains of hypoxic rats (Fig. 5, j-k). Histological sections from control rats show more NF-H-positive neurons, stronger immunoreactivity of the entire soma and neurites, more elaborated apical and basal dendrites, and prominent neuropil in layers II-VI. Based on neurofilament analysis, control brains exhibit a more advanced development of the NF-H-positive neuronal population (Figures 5, f-g, arrows). On the other hand, hypoxic rats exhibit fewer cells, weaker NF-H expression, smaller soma, stained only the proximal segment of the apical dendrites, as well as less complex neurite arborization and neuropil (Fig. 5, j-k, arrows). Given the preserved gross cytoarchitecture on Nissl staining, these findings are interpreted as evidence of altered cytoskeletal maturation rather than neuronal loss.

Conversely, the PV immunoreactivity displayed upregulated expression of PV protein in the neuropil and soma of interneurons in the cortex of the hypoxic group. PV staining revealed the qualitative and quantitative difference in the PV-positive cells in the ACC (Fig. 6). The observed upregulated diffuse staining of the neuropil in the hypoxic group and in the soma and proximal dendrites of the hypoxic rats is shown in Fig. 6, f, arrow. An indicatively higher number of PV-positive interneurons was counted in all layers of the ACC of hypoxic rats (Welch’s t-test, p = 0.137, Fig. 6, g), with the highest difference observed in layer V (Welch’s t-test, p = 0.061, Fig. 6, i).

**Fig. 6.**
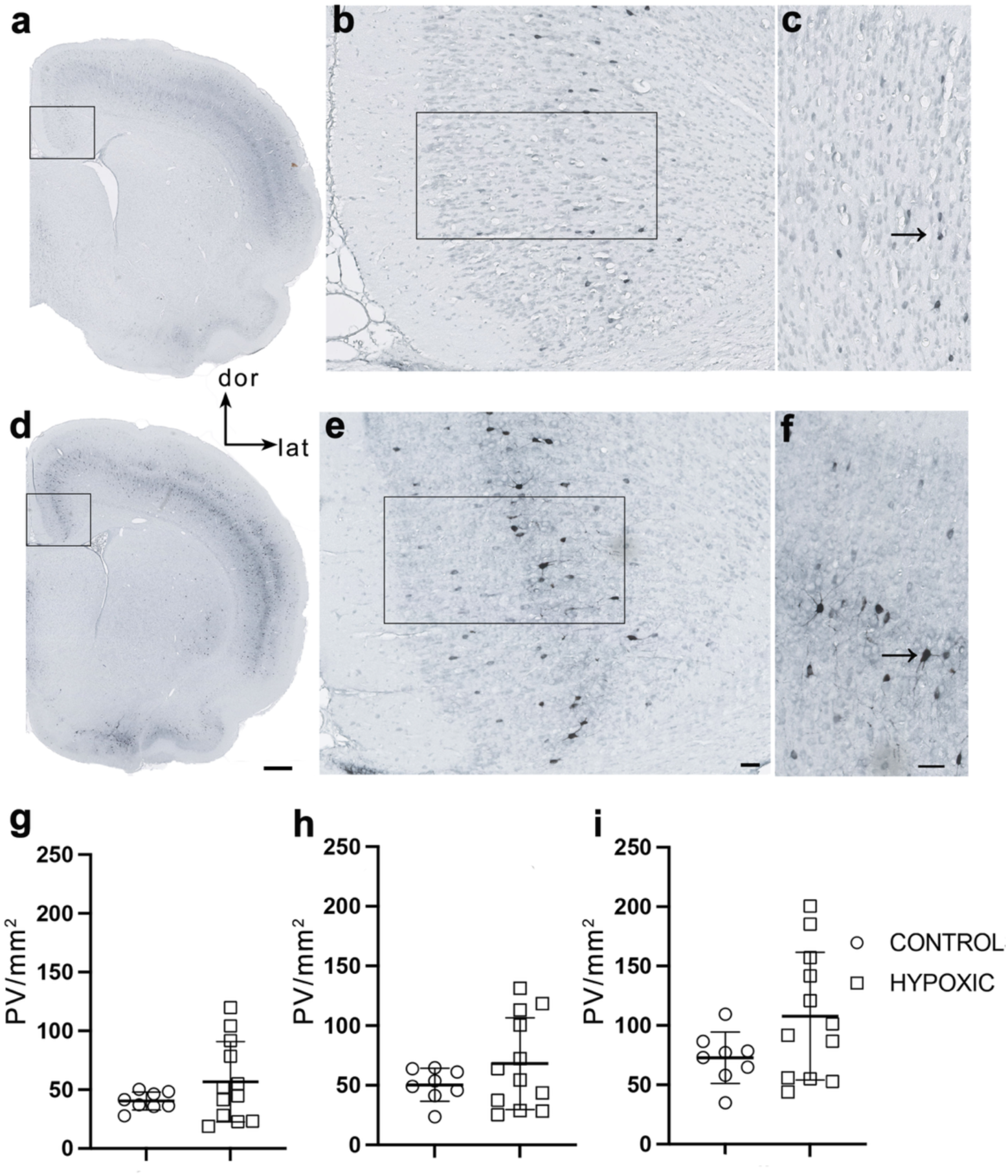
Accelerated maturation of PV neurons in the ACC following MPH. Coronal sections at the level of the ACC (∼+1.28 mm from Bregma) stained for PV in control (a–c) and hypoxic rats (d–f). Black rectangles in (a, b; d, e) indicate regions enlarged in (b, c; e, f). Hypoxic rats showed increased diffuse PV staining (d–f), most prominently in layer V of the ACC (f), compared to controls (a–c). Stronger PV immunoreactivity is observed in the soma and proximal dendrites of hypoxic rats (arrows in f) compared to controls (arrow in c). Quantification indicates an increased number of PV-positive interneurons per mm² in the ACC (g; Welch’s t-test, p = 0.137), predominantly in infragranular layers (h; Welch’s t-test, p = 0.164), with the most prominent difference in layer V (i; Welch’s t-test, p = 0.061). Data are presented as mean ± SD. Scale bars: 0.5 mm in (d); 50 μm in (e, f), applicable to (b–c)

A statistically significant increase in PV-positive interneuron number was observed in the dorsolateral cortex, the sensory neocortex (Fig. 7). The dorsolateral cortex in the hypoxic group showed narrow and less distributed diffuse staining, more evident for the supragranular layers (Fig.7, e), compared to controls (Fig.7, b). The statistically significant effect of perinatal hypoxia on PV neuron number is observed in somatosensory/sensory cortex (, p<0.001), predominantly in infragranular layers (Welch’s t-test, P=0.002). The most remarkable difference between groups was observed again in layer V (arrows), with a higher number of PV/mm^2^ in hypoxic group (Welch’s t-test, p< 0.001) (Fig. 7, i). The quantitative analysis aligns with the qualitatively observed prominent difference in layer V. Moreover, stronger PV expression is observed in the soma, and proximal dendrite segments of the sensory cortex in hypoxic rats (Fig.7, f).

**Fig. 7.**
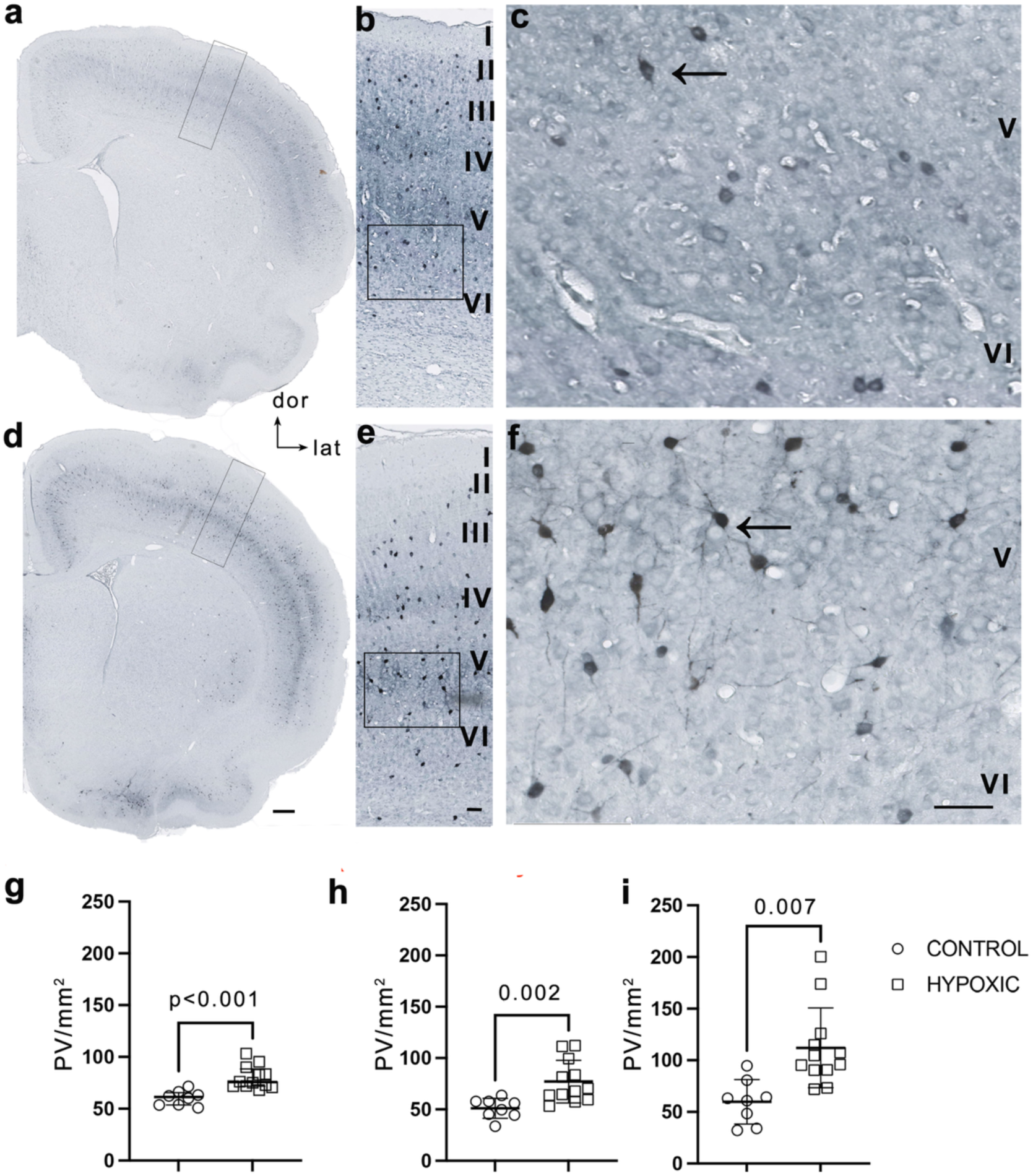
Increased number and accelerated maturation of PV neurons in the S1 cortex at P15 following MPH. Coronal sections at the level of the sensory cortex (∼+1.28 mm from Bregma) stained for PV protein in control (a–c) and hypoxic rats (d–f). Black rectangles in (a, b; d, e) indicate regions enlarged in (b, c; e, f). PV immunostaining in the primary somatosensory (S1) cortex showed less distributed diffuse staining in hypoxic rats (d–f), with the reduction most prominent in the supragranular layers (e, f). Qualitative analysis revealed stronger PV immunoreactivity in the soma and proximal segments of apical and basal dendrites of hypoxic rats (arrows in e, f), compared to a more immature staining pattern in controls (arrows in b, c). Quantification of PV-positive neurons per mm² (j–l) showed a significant effect of MPH in S1 (j; Welch’s t-test, p < 0.001). A higher number of PV interneurons in hypoxic rats is observed in infragranular layers (k; Welch’s t-test, p = 0.002). The largest difference was observed in layer V of S1 (l), with a significant increase in PV cell numbers in hypoxic rats (Welch’s t-test, p = 0.001). Data are presented as mean ± SD. Scale bars: 0.5 mm in (d) applies to (a); 50 μm in (e, f) applies to (b, c)

The quantitative analysis of Cplx3 expression in the SpN (VIb layer), a neuronal population critically important for development, showed no differences in its number after MPH. Nevertheless, we observed a qualitative difference in the neuronal phenotype revealed by Cplx3 staining, specifically, upregulation of Cplx3 expression in all SpN neurons in hypoxic rats compared to variably stained SpN in controls (Supplementary Fig. S4).

The qualitative analysis of the overall expression of hyaluronan and proteoglycans’ glycosaminoglycans (GAG) visualized by Mowry staining (Supplementary Figure S5, a, c), as well as the expression of proteoglycans, including keratan sulfate (Supplementary Figure S5, b, d), neurocan, versican, and lumican (not shown), did not consistently differ between controls and hypoxic rats. This study did not reveal significant quantitative differences in the amount of the expressed GAG component of the ECM during early postnatal development up to P15, assessed indirectly by measuring the intensity of Mowry staining (Supplementary Figure 5e). However, analysis of WFA labeling revealed qualitative differences in the staining pattern of diffusely distributed ECM, most prominently in the infragranular layers of the ACC and dorsolateral cortex (Supplementary Figure S6, a-c and e-g), indicating changes in ECM glycosylation pattern after MPH. In the brains of control rats, WFA immunoreactivity was evenly distributed across cortical layers, reflecting cortical lamination (Supplementary Figure S6, a-c), whereas in hypoxic rats after MPH, WFA staining was most prominent in the infragranular layers. Only immature, reticular, forerunners of jet undeveloped perineuronal nets (PNNs) were observed in both groups (Supplementary Figure S6, d and h)

The immunohistochemical staining and qualitative analysis of MBP revealed lower expression of MBP-positive oligodendrocytes indicative of a delay in fiber myelination in ACC and the dorsolateral (around Bregma level 0.48; Paxinos and Watson 2014) neocortex in the juvenile hypoxic rats (P15) (Fig. 8). Compromised myelination was observed in the lagging progression of myelination from the deeper to the superficial parts of the cortical layers in the hypoxic group, visible in the ACC, as well as in the dorsolateral somatosensory cortex (Fig. 8, d, g). Compared to controls, hypoxic rats had an unmyelinated layer V and only a narrow zone of myelinated fibers in layer 6, medial to the genu of the corpus callosum in ACC (Fig. 8d).

**Fig. 8.**
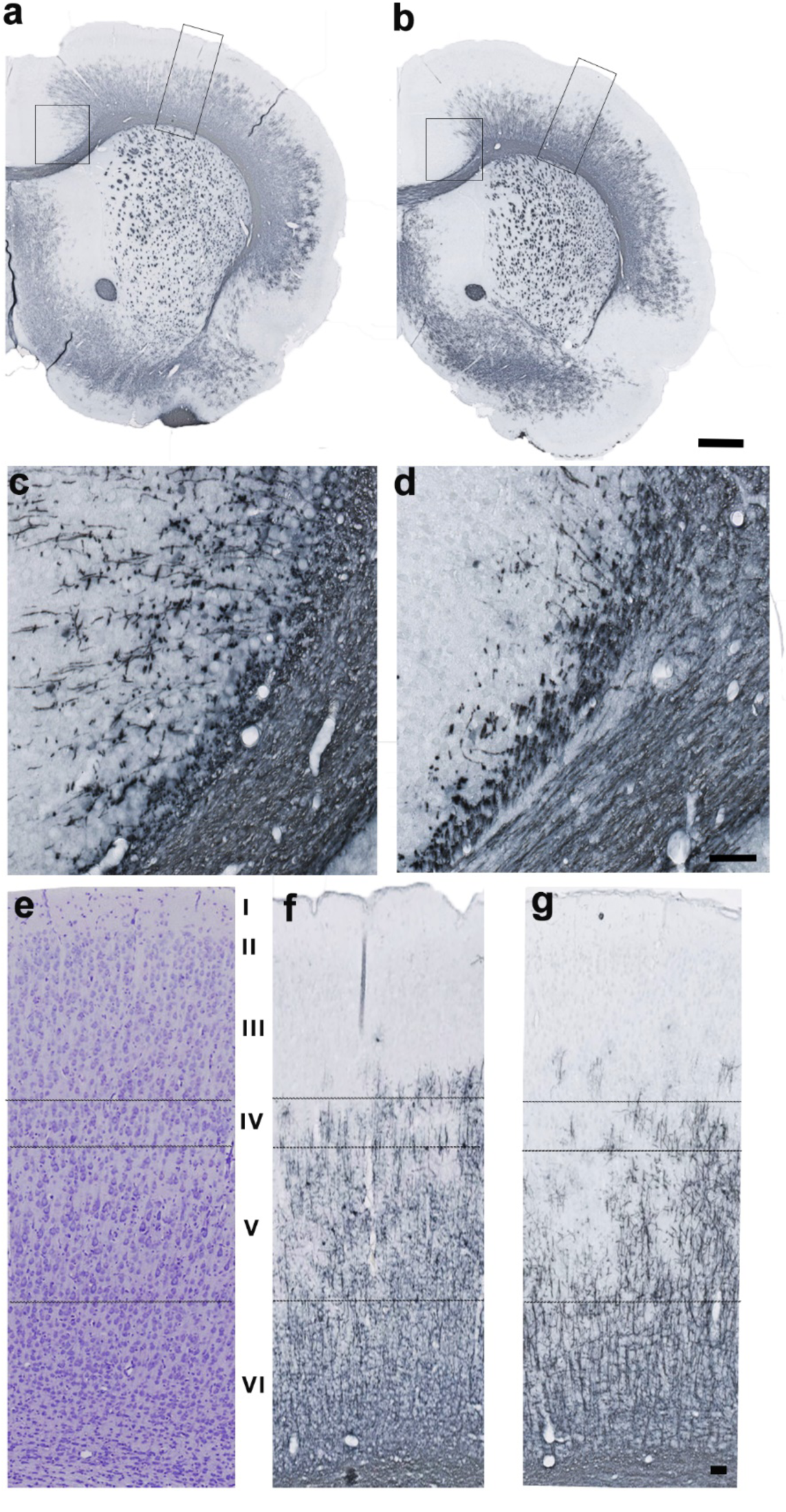
Disrupted myelination in the ACC and S1 cortex at P15 following MPH. Coronal sections at the level of the ACC (∼+0.48 mm from Bregma) stained for MBP in control (a, c, f) and hypoxic rats (b, d, g). MBP immunostaining revealed myelination deficits in the ACC and primary somatosensory cortex (S1) in hypoxic rats. In the ACC, a reduction of myelinated fibers was evident in cortical layers VI and V, with absence of fibers in layer V and only a narrow, myelinated band in layer VI medial to the genu of the corpus callosum (d), compared to controls (c). Similarly, in S1, hypoxic rats displayed reduced myelination, particularly in layers IV and V (g), compared to controls (f). Scale bars: 0.5 mm (b); 50 μm (d, g). Identical magnification is shown for panels (a-b), (c–d), and (e-g)

## DISCUSSION

This study revealed the consequences of MPH on cerebral cortex development in rats, demonstrating that even a single moderate hypoxic episode leads to significant structural and functional alterations in brain organization. Specifically, assessment of neurodevelopmental milestones showed reduced performance intensity and delayed onset and cessation of reflexes, without locomotor deficits, indicating impairment and therefore greater vulnerability of developing somatosensory circuits. At the structural level, the most pronounced effects were detected in the ACC by in vivo MRI, histologically characterized by changes in the dendritic organization of principal neurons and an increased number of PV-expressing interneurons.

### Differential vulnerability of early developmental reflexes

Differential CNS vulnerability, as assessed behaviorally, identified the brainstem, somatosensory, and sensorimotor connectivity as most vulnerable to MPH in rats. After MPH, rats exhibited a delayed onset and prolonged persistence of key reflexes, including the vibrissae placing and rooting reflexes, indicating disrupted sensory input processing during a critical developmental window.

The maturation of neurological reflexes represents a hallmark of nervous system development, while delays in their suppression or persistence beyond the expected developmental window indicates neurodevelopmental impairment (Fox 1964; Dobbing and Smart 1971; Altman and Sudarshan 1975; Donatelle 1977). For example, retained primitive reflexes in children with autism spectrum disorder (ASD) have been related to functional brain dysconnectivity due to maturational delays of sensory, motor, and emotional regulation (Melillo et al. 2022). Perinatal factors, including neonatal hypoxia–ischemia, have been reported to influence these reflexes (Lubics et al. 2005). Hypoxia activates stress-responsive pathways, including HIF-1α signaling and catecholaminergic systems, which can transiently increase arousal and locomotor activity (Schönenberger and Kovacs 2015). In our study, following MPH, the rooting and vibrissae placement reflexes were significantly less pronounced, suggesting lesions of the brainstem and/or sensorimotor processing or connectivity and a consequent delay in the maturation of the early sensory-motor circuit. However, a rapid recovery of these circuits, with functional restoration within less than two weeks, was observed. Although hypoxic rats outperform controls in the surface righting test, this finding may not necessarily reflect accelerated maturation of the motor circuits. The hyperactivity could reduce righting latency without indicating genuine neurodevelopmental improvement. Moreover, early compensatory responses to hypoxia are often followed by delayed deficits in synaptic organization, myelination, and motor coordination. Therefore, the superior performance observed at P5–P6 in hypoxic rats may represent a short-lived adaptive or stress-induced phenomenon rather than a lasting functional benefit. This interpretation aligns with recently published findings of persistent behavioral alterations and imbalances in mesothalamic dopaminergic signaling later in development (P33-P43) following MPH, suggesting that early reflex enhancements are masking long-term neurochemical dysregulation rather than representing developmental acceleration (Nikolic et al. 2023).

### Principal neurons’ developmental delay and cytoarchitecture remodeling

Previous studies investigating the effects of generalized mild-to-moderate perinatal hypoxia in the rat brain (Trnski et al., 2022; Nikolic et al., 2023) found no evidence of acute or chronic reactive gliosis or its sequelae in mid-adolescent (P35 - 45) or adult (>P60) rats, thereby directing the present study toward neuronal and ECM components.

The radial arrangement of apical dendrites of cortical neurons is well described in humans and rodents as a part of the cortical structural radial units of the developing mammalian cortex (Gabbott and Bacon 1996; Peters et al. 1997; Lewis and Van Essen 2000; Rockland and Ichinohe 2004; Buxhoeveden et al.). The neocortex shows prolonged maturation, especially in the associative and “prefrontal” medial cortical regions (Ulyngs et al. 2003; Vogt and Paxinos 2014; Pallomero-Gallagher and Zilles 2015; Preuss and Weiss 2022). Finally, in the mature neocortex, vertical dendritic bundles are also masked by extensive dendritic arborization of basal dendrites and neuropil elaboration (Innocenti and Vercelli 2010). MAP2 is an excellent molecular marker of these organizational units and their maturation sequence. The presence of prominent, robust, vertically oriented, bundled MAP2-positive apical dendrites in ACC layers II, III, and V in hypoxic rats confirms the altered development of cortical cytoarchitecture. We found increased MAP2 expression in dendrites following MPH, in contrast to the reduced MAP2 expression and neuronal loss observed after severe ischemic brain injury in rodents (Dawson and Hallenbeck 1996; Mages et al. 2021) and humans (Kühn et al. 2005). The NF-H staining further confirms the disrupted neuronal organization and maturation in the neocortex. In hypoxic rats, specific NF-H-positive neurons, primarily large pyramidal long-projection neurons in layers III and V, exhibited underdeveloped neurite morphology and modest neuropil, especially pronounced in the ACC. In a mouse model of chronic hypoxia, similar alterations in this neuronal population were observed, including disrupted dendritic arborization of pyramidal neurons and disorganization of cortical neuropil (Fagel et al., 2006).

### Predominant advanced differentiation of interneurons and inhibitory circuits

PV-positive interneurons exhibit high metabolic demands due to their fast-spiking activity, rendering them particularly vulnerable to oxidative stress and energy deficits under hypoxic conditions (Kann 2016; Fowke et al. 2018). In this study, we found that the observed upregulation of PV in the neocortex of adult rats following perinatal hypoxia (Trnski et al. 2022), occurs early in development, and is present already by P15. It is known that early-developing interneurons exert excitatory, depolarizing GABAergic activity, which later shifts to hyperpolarizing, inhibitory GABAergic activity, characteristic of hyperpolarizing inhibitory GABAergic interneurons (Luhmann and Prince 1991). This GABAergic shift occurs during the second postnatal week in rodents (Ben-Ari 2002). In the rat neocortex, these GABAergic synapses gradually mature during the first postnatal month (Luhmann and Prince 1991), influencing proliferation (Haydar et al. 2000), cell migration (Luhmann et al. 2015), dendritic development (Sernagor et al. 2010), and synaptogenesis (Wang and Kriegstein 2008). The observed changes in NF-H expression may also reflect the effects of insufficient depolarizing GABA input and, consequently, impaired dendritic outgrowth and delay in pyramidal neuron differentiation. Thus, the increased number of PV-interneurons in the cingulate and somatosensory cortices after MPH indicates premature maturation of inhibitory networks, which could first affect developmental processes and later affect excitatory-inhibitory homeostasis, contributing to the observed behavioral aberrations as well. These findings underscore the vulnerability of interneurons in the critical GABAergic shift, leading to precocious maturation of the inhibitory network (Josh Huang et al. 1999; Fagiolini and Hensch 2000; Maffei et al. 2006). With respect to preventing excessive neuronal excitation through GABA inhibition, accelerated maturation of these neurons may also represent a plasticity response to hypoxia (Ivanov et al. 2015).

### Persistence and plasticity response of transient developmental circuitries

Available comparative and developmental studies suggest that the human fetal/perinatal cerebral cortex exhibits more complex and prolonged expression of ECM constituents and more prominent ECM-rich compartments than the rodent cortex (Jovanov Milosevic et al., 2014; Kostovic et.al., 2014a; Fietz et al., 2012; Amin and Borrell, 2020; Long and Huttner, 2022). This is particularly evident by the abundance of GAGs and proteoglycans in the marginal and subplate zones, and in outer radial glia niches (Jovanov Milosevic et al., 2014; Kostovic et al., 2014a; Fietz et al., 2012; Pollen et al., 2015; Long et al., 2018). These interspecies differences may explain why we were unable to detect significant changes in most ECM components in the rat brain after MPH at P15 using our methodology.

At the same time, our results showed a more immature appearance of the diffuse WFA staining pattern in hypoxic rats, suggesting dysregulation of cortical ECM glycosylation (at least the lectin-binding component) and possibly a delayed or disrupted transition from diffuse to condensed expression, consequently altering PNN formation and maturation. This effect most likely reflects the early occurrence of alterations in the proteoglycan components of the ECM, previously reported in the adult rat brain following MPH (Trnski et al. 2022). For some neuronal circuits, maturation and critical period closure are determined by ECM transitions from a diffuse distribution of proteoglycans around to highly organized lattice-like structures surrounding neurons (PNNs) (Wang and Fawcet, 2012). In addition, injury-induced changes in glycosylation and sulfation patterns of glycosaminoglycans (Alonge et al., 2021) could modify lectin-binding properties without necessarily altering core protein proteoglycan expression, providing another plausible explanation for the different diffuse WFA pattern observed in the hypoxic cortex.

Current evidence underscores that SpNs are particularly vulnerable to neonatal hypoxia in rodents (McQuillen et al. 2003; Mikhailova et al. 2017; Sheikh et al. 2019), and their disruption may impair early cortical network formation, leading to long-term alterations in connectivity (Kanold 2009). SpNs are among the earliest-maturing cortical neurons in rodents and serve as transient synaptic targets that guide thalamocortical afferents, linking thalamic input to cortical circuits (Molnár et al. 2020). Most SpNs undergo programmed cell death; in rodents, apoptosis peaks in the first postnatal week and is largely completed within the first two weeks (Al-Ghoul and Miller 1989; Ferrer et al. 1990; McQuillen et al. 2002; McQuillen and Ferriero 2005). In the present study, sustained upregulation of Cplx3 suggests a developmental delay and may reflect compensatory plasticity.

In humans, peak subplate development coincides with the period of highest susceptibility to periventricular leukomalacia (PVL), around 24 gestational weeks (McQuillen et al. 2003; Kostović et al. 2014a; Kostović et al. 2014b). By forming the first functional cortical networks, they mediate early excitation between the thalamus and neocortical layer IV, and their injury has been implicated in multiple neurodevelopmental disorders (Rocha-Ferreira and Hristova 2016). Following thalamocortical circuit refinement after perinatal injury, most SpNs are eliminated through programmed cell death (McQuillen and Ferriero 2005). Notably, the prolonged persistence of SpNs in the human prefrontal cortex suggests a role in extended cortical maturation and sustained plasticity until synaptic stabilization is achieved (Van Eden et al. 1991). In this context, the observed qualitative changes in SpN phenotype, together with prolonged Cplx3 expression, likely reflect delayed maturation and a compensatory plasticity response to MPH-induced injury in rats.

### Detectability of MPH effects by in vivo MRI

Although ultrasonography remains the primary bedside method for monitoring the neonatal brain (Hüppi 2004), it still has limited diagnostic value for diffuse brain injuries. By contrast, MRI is ideally suited for assessing diffuse and microstructural alterations in neuronal organization, myelination, and white matter integrity (Hüppi 2004). Diffusion tensor imaging (DTI) is sensitive to tissue microstructure by quantifying the magnitude and directionality of water diffusion (Song et al. 2003; Tao et al. 2012). Thus, DTI-derived parameters provide indirect imaging biomarkers of white matter integrity and dendritic organization (Beaulieu 2002). Given this, we performed whole-brain voxel-wise DTI MRI and volumetric analyses to identify the most affected, and hence most vulnerable, cortical regions by this MPH injury. Collectively, MRI findings suggest that the single MPH event provoked a microstructural tissue rearrangement with symmetrical distribution, indicating the vulnerability of specific brain regions rather than a generalized injury. The most prominent changes were in the cerebral cortex, most notably in the ACC, suggesting heightened vulnerability of this region, followed by the dorsolateral and sensory cortices. This pattern may reflect the regional susceptibility of parasagittal cortical areas known as vascular watershed zones between the anterior (ACA) and middle cerebral artery (MCA) territories, as in the perinatal human brain, where these border-zone regions are particularly vulnerable to hypoperfusion and are commonly involved in partial-prolonged hypoxic–ischemic injury (Volpe 2008; de Vries and Groenendaal 2010; Douglas-Escobar and Weiss 2015; Misser et al. 2020; Bobba et al. 2023). In the rat brain, the ACA–MCA watershed region encompasses the parasagittal medial cortex, including the ACC, prelimbic cortex, and infralimbic cortex, as well as medial portions of the motor and somatosensory cortices where perfusion from these arterial territories overlaps. This region lacks sharp anatomical boundaries and is instead defined functionally by transitional perfusion between the anterior cerebral artery and middle cerebral artery territories, as demonstrated by vascular mapping and ischemia models (Coyle 1975; Tamura et al. 1981; Longa et al. 1989).

At P15, after MPH, the immature ACC exhibits a significant increase in FA, accompanied by a non-significant but detectable increase in AD and a slight decrease in RD, suggesting fine microstructural changes in the cytoarchitecture of this area, such as compromised neuronal fiber integrity and organization. Disrupted neuronal development, characterized by fewer and shorter dendritic branches, presents fewer pathways for water to diffuse parallel to the pial surface (Leigland et al. 2013). Accordingly, MAP2 and NF-H staining suggest that the elevated FA values may correspond to hypoxia-induced, less elaborated dendritic trees in the developing cortex and tighter packing of neuronal fibers (dendrites and axons), which together contribute to altered cortical fiber organization. Considering the timing of MR imaging after MPH, and developmental stage, these findings in ACC are most likely residual to the initial cytotoxic damage of axons, consequent reduction of extracellular space (resulting in restricted RD and increased FA), and simplified axonal and dendritic arborization in these regions, which is in line with the behavioral developmental delay observed in hypoxic rats. In healthy animals, FA values decrease as neurons differentiate and the cortex matures (Neil et al. 1998; Bockhorst et al. 2008; Ball et al. 2013; Breu et al. 2019). Therefore, we propose that FA changes reflect developmental delay, simplified neurite complexity, and prolonged ACC maturation in hypoxic rats. These findings are comparable to the delayed brain maturation observed in neonates who suffered from mild hypoxic-ischemic encephalopathy (Gao et al. 2015).

Although white matter is less developed in rodents’ brains than in humans’, it undergoes rapid postnatal maturation characterized by increases in axonal diameter, oligodendrocyte number, and myelination (Morken et al. 2013). MBP expression did not reveal significant differences in myelination in the cortical layers of ACC area, where increased FA was observed. Thus, we suggest that the change in FA in the cortical gray matter of juvenile rats following MPH likely reflects cytoarchitectural differences rather than compromised myelination. These findings are consistent with the previously reported relationship between MR anisotropy in the cerebral cortex and features of unmyelinated tissue (Reveley et al. 2022). The prevailing view is that delayed myelination is most likely due to the vulnerability of pre-oligodendrocytes and their progenitors to early hypoxia (Liu et al. 2001) on the one hand, and to axonal subacute post-hypoxia reduction (accompanied by partial recovery and reorganization) on the other. In hypoxic rats, DTI showed elevated AD in the upper cortical layers of posterior sensory cortices, while RD and FA remained unchanged. In parallel, MBP expression was reduced to the lower cortical layers. Thus, the differences in myelination between groups, in the ACC and DL at P15, are less likely to be detected by DTI in our study. This spatial dissociation results from layer-specific organization and, consequently, lesions specific to certain layers following perinatal hypoxia. The lower cortical layers (V/VI), which give rise to early-myelinating corticofugal projections (e.g., corticospinal and corticothalamic pathways), exhibit delayed or impaired myelination. In contrast, increased AD in the upper layers (II/III) reflects reduced dendritic complexity or sparse axonal terminations due to affected afferent input and intracortical connectivity. This combination confirms the diverse and lasting neurodevelopmental impact of MPH, with both myelin and axonal structures affected.

The vulnerable regions identified in this MPH rat model correspond to those reported in the human preterm brain, supporting the model’s translational relevance (Volpe 2009; Back and Miller 2014). The volume of the inferior colliculi was significantly increased, and a trend toward increased volume in the lateral thalamic nuclei was observed, yet these regions did not show differences in diffusivity parameters. This is consistent with reports of human preterm studies claiming that DTI metrics may remain unchanged despite ongoing developmental alterations (Hüppi and Dubois 2006; Counsell et al. 2008). In the rat model, two weeks after MPH, regions were not enlarged due to swelling (neither cellular nor extracellular) but rather most likely due to a delay in developmental pruning or to excessive synaptogenesis, with exuberant ECM and growth factors, as a post-lesion plasticity response (McClendon et al. 2014). This pattern aligns with the concept of dysmaturation rather than overt structural injury (Smyser et al. 2010; Back and Miller 2014) and correlates well with the behavioral aberrations observed in rats after MPH.

## Conclusion

We provide evidence that a single episode of MPH is sufficient to trigger measurable, early (P15) reorganization of cortical circuitry that underlies the long-lasting effects observed in mature brains in this hypoxia model (Trnski et al. 2022; Nikolic et al., 2023). The ACC emerges as the most vulnerable region to generalized MPH, indicating region-specific susceptibility during this developmental window. At the cellular level, MPH appears to disrupt the maturation trajectory of pyramidal neurons, as evidenced by reduced dendritic complexity and altered cytoskeletal organization, consistent with impaired establishment of excitatory connectivity. In parallel, an increased number of PV-interneurons and neuropil elaboration, indicating precocious differentiation of inhibitory networks, likely reflects activity-dependent compensatory mechanisms. Transient SpNs, rather than being eliminated, exhibit prolonged persistence, suggesting altered specification, tuning, and pruning, thereby extending cortical circuit formation. We propose that MPH differently shifts the developmental timing of cortical circuit assembly by delaying excitatory maturation, accelerating inhibitory differentiation, and prolonging persistence of SpNs thereby destabilizing the excitation–inhibition balance and impairing network refinement. These convergent alterations lead to aberrant circuit integration and are associated with early behavioral deficits and increased FA in the ACC, linking microstructural changes to the MRI signature of MPH. Together, the data define a mechanistic framework in which subtle hypoxic insults perturb developmental trajectories of cortical networks, establishing critical windows of vulnerability and potential cellular and molecular research targets for timely intervention.

## Acknowledgments

The authors thank Academician Ivica Kostović and Professor Nenad Bogdanović for critically reviewing the final manuscript and providing valuable suggestions for its improvement. They also thank Maja Horvat Božić and Božica Popović for their excellent technical support, as well as all colleagues at the Laboratory for Regenerative Neuroscience – GlowLab (RRID:SCR_022701) for their excellence in animal MRI.

## STATEMENTS & DECLARATIONS

### Author Contributions

Material preparation, data collection, and analysis were performed by M. DC., S. TL., S. S., D. dC., M. BR., and E. K. The first draft of the manuscript was written by M. DC. and S. TL. Statistical analysis was performed by A. Š. Supervision of the formal analysis, critical data selection, literature review, and manuscript editing was carried out by M. DC., S. TL., S. S., D. dC., E. K., M. BR., I. K., A. Š., K. I., D. C., M. J., and N. J. M. The study design and conceptualization by N. J. M. All authors commented on previous versions of the manuscript and read and approved the final manuscript.

### Data Availability

The datasets generated during the current study are available on request.

### Ethics approval

All animal experiments complied with Croatian regulations for experimentation on animals (NN 102/2017 and 32/19; NN 55/2013; 39/17 and 116/2019). The study design and experiments wereapproved by the Ethics committee of the University of Zagreb and the national Ethics and Animal welfare bodies (EP231/2019; UP/I-322-01/19.01.75).

