## Supplementary material for "Mild perinatal hypoxia uncouples excitatory–inhibitory circuit maturation and reprograms neocortical organization": Supplementary_Figures_bioRxiv.pdf

-

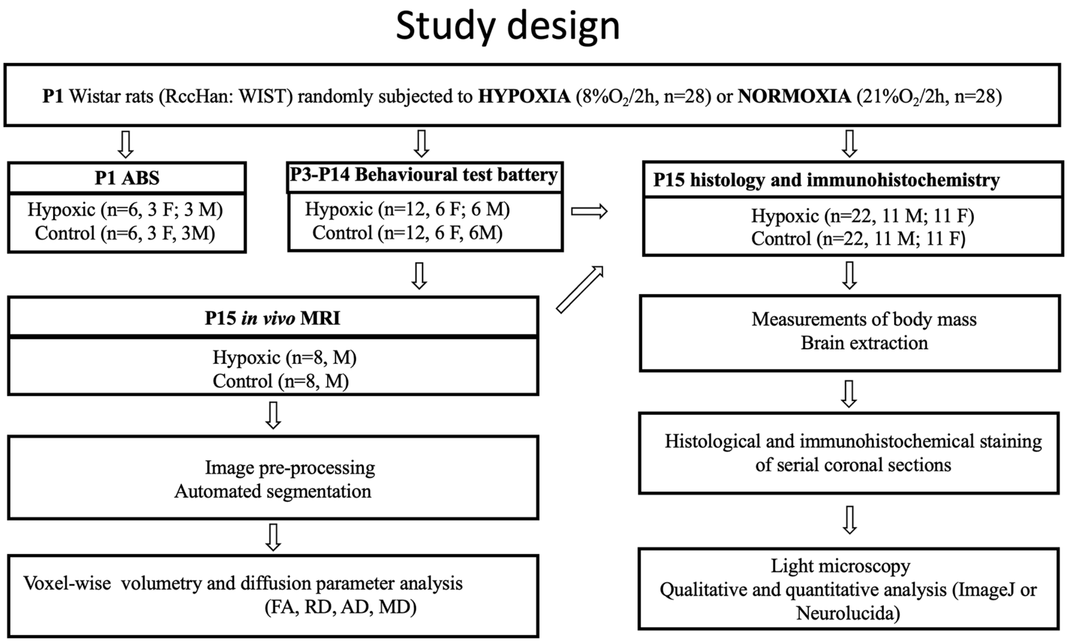


**Supplementary Fig. S1 Study design**

Overview of the experimental design and group allocation. A total of 56 rats of both sexes were included: 28 exposed to hypoxia (8% $O_2$ for 2 h) and 28 controls (room air for 2 h). Immediately after exposure, 12 rats (n = 6 per group) were used to assess acid–base status (ABS). The remaining 44 animals were sacrificed at P15 for histological and immunohistochemical analyses (antibodies listed in Supplementary Table 3). Of these, 24 rats (12 hypoxic, 12 control) underwent behavioral testing from P3 to P14. A subset of males (n = 16; 8 per group) underwent in vivo MRI at P15.

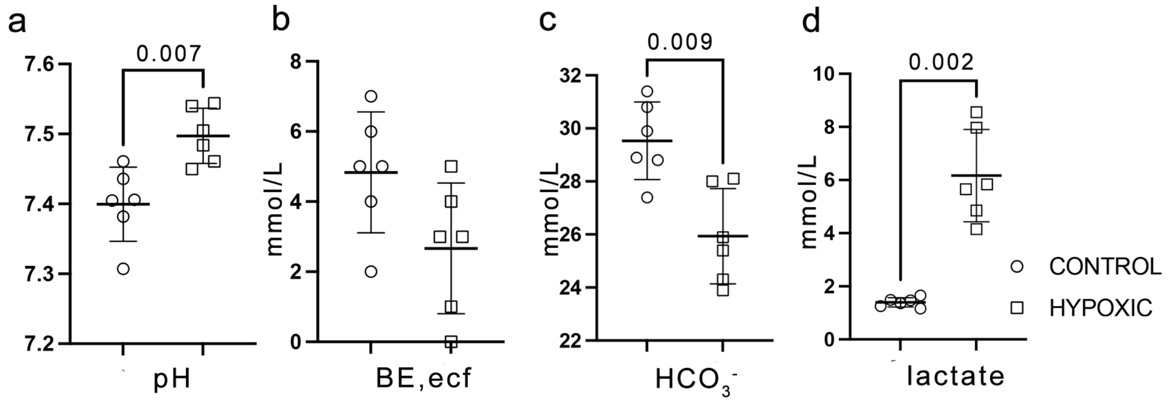


**Supplementary Fig. S2 Acid–base status (ABS)**

Acute response to hypoxic exposure was assessed by measuring (a) pH, (b) base excess in extracellular fluid (BE_ecf), (c) $HCO_3^-$ (bicarbonate), and (d) lactate levels. Tissue hypoxia was confirmed by increased lactate levels (d; p = 0.002, Mann–Whitney test). Compensatory response was indicated by increased pH (a; p= 0.007, Mann-Whitney test) and decreased $HCO_3^-$ levels (c; p=0.009, Mann-Whitney test) in hypoxic rats. Data are presented as medians with interquartile ranges (IQRs).

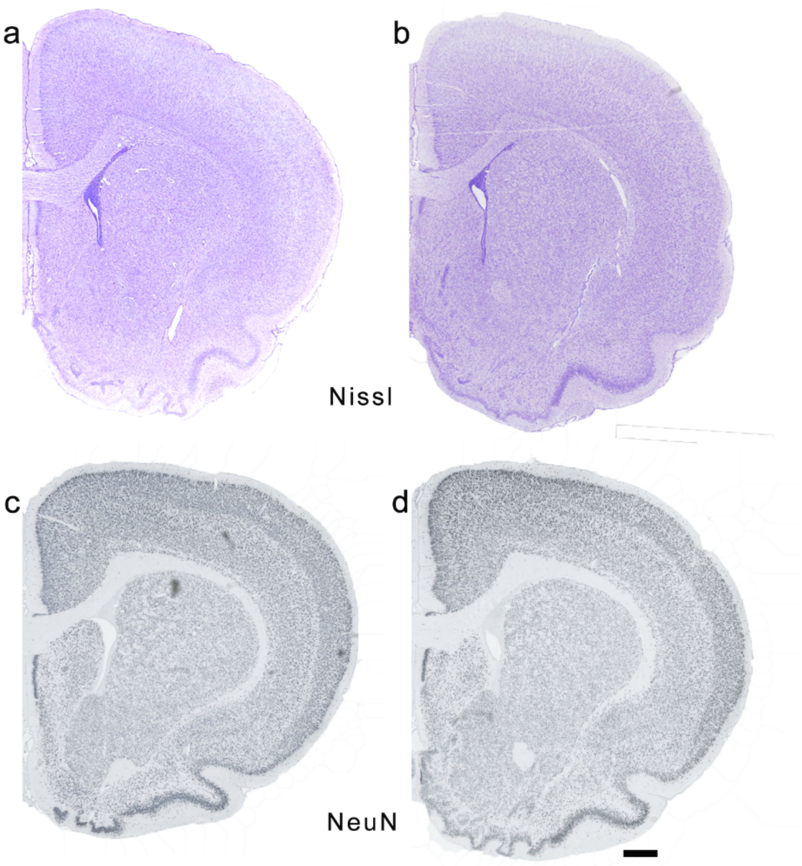


**Supplementary Fig. S3 Nissl and NeuN staining show no gross histopathological alterations or neuronal loss following MPH**

Coronal sections at ~+2.28 mm from Bregma were stained with Nissl (a, b), NeuN (c, d). Nissl staining shows preserved morphology and intact cortical lamination in hypoxic rats (a), comparable to controls (b). The NeuN immunostaining, by qualitative analysis, shows no evidence of neuronal loss or altered expression in hypoxic rats (c) compared to controls (d).

Scale bar: 0.5 mm in (d) also applies to (a-c)

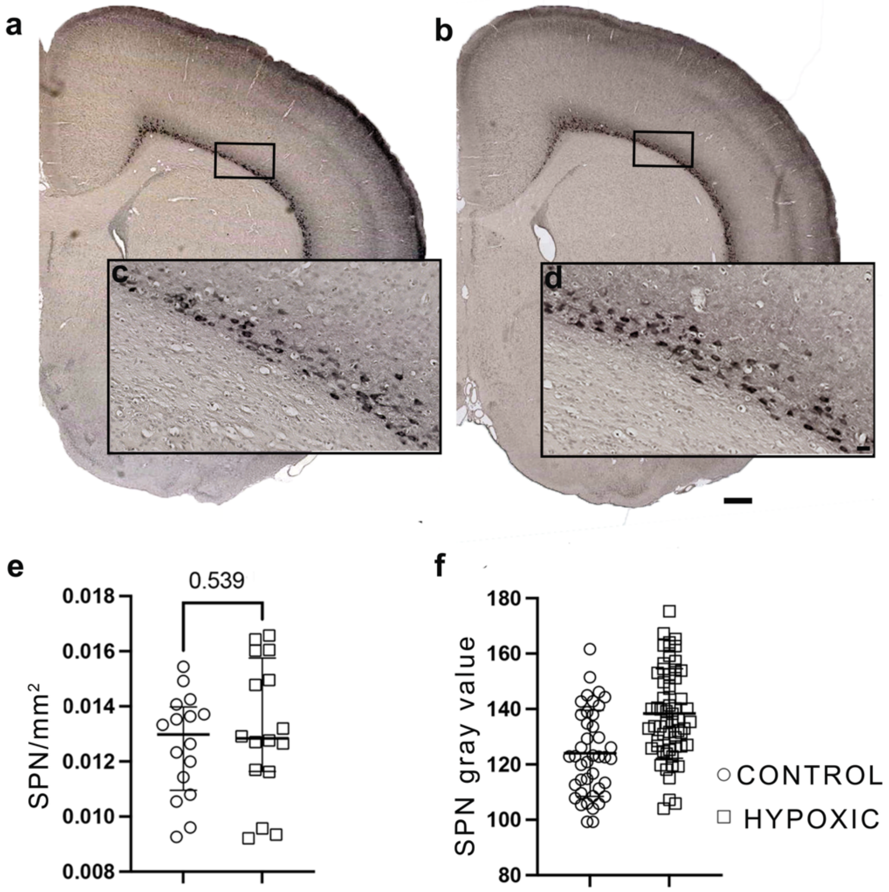


**Supplementary Fig. S4 Cplx3 expression shows unchanged subplate neuron number density and increased and prolonged Cplx3 protein expression at P15 following MPH**

Coronal sections at ~+2.28 mm from Bregma stained for the subplate neuron (SpN)-specific marker Cplx3 in control (a) and hypoxic rats (b). Black boxes in (a, b) indicate regions enlarged in higher-magnification images (c-d). Increased cytoplasmic Cplx3 immunoreactivity is observed in SPNs of hypoxic rats (b) compared to controls (a).

Quantification of staining intensity (f) shows a significant increase in Cplx3 signal in hypoxic rats (Welch's t-test, $p < 0.0001$). Individual gray values (0–255 scale, higher values indicating darker pixels) are presented as mean ± SD. In contrast, SpN counts (e) show no difference between groups (median ± IQR, $p = 0.539$; Mann–Whitney test), indicating preserved SpN density following mild perinatal hypoxia.

Scale bars: 0.5 mm in (b) applies to (a), 25 µm in (d) also applies to (c)

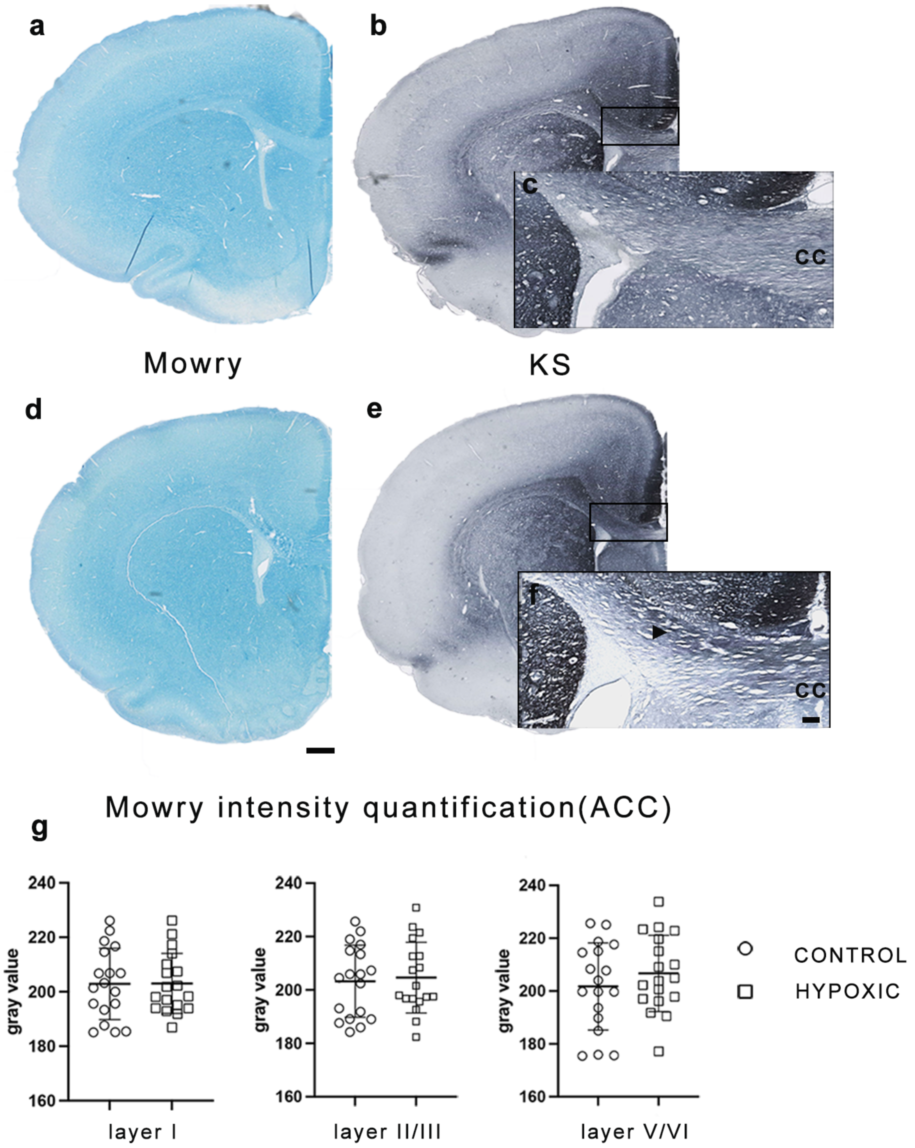


**Supplementary Fig. S5 Hyaluronan and keratan sulfate expression in the ACC of control and hypoxic rats at P15**

Coronal sections at the level of the anterior cingulate cortex (ACC; ~+2.28 mm from Bregma) from control (a–c) and hypoxic rats (d–f), stained with (a, d) colloidal iron (Mowry), (b, e) anti-keratan sulfate (KS). Mowry staining shows no difference in hyaluronan (HA) expression in the ACC of hypoxic rats (d) compared to controls (a). Quantification of Mowry staining intensity (g; Welch's t-test) shows no difference in HA across cortical layers I–VI. Analysis was performed in three regions of interest (ROIs): layer I, layers II–III, and layers V–VI; data are presented as mean ± SD of gray intensity values.

KS-positive diffuse ECM is present in the ACC of both groups (b, e), with the strongest labeling in layers I and III. Black boxes in (b, e) indicate regions enlarged in higher magnification. Dorsal callosal fibers show stronger KS immunoreactivity in hypoxic rats (arrowhead in f) compared to controls (c).

Scale bar: 0.5 mm in (d) applies to (a–e); 50 µm in (f) applies to (c)

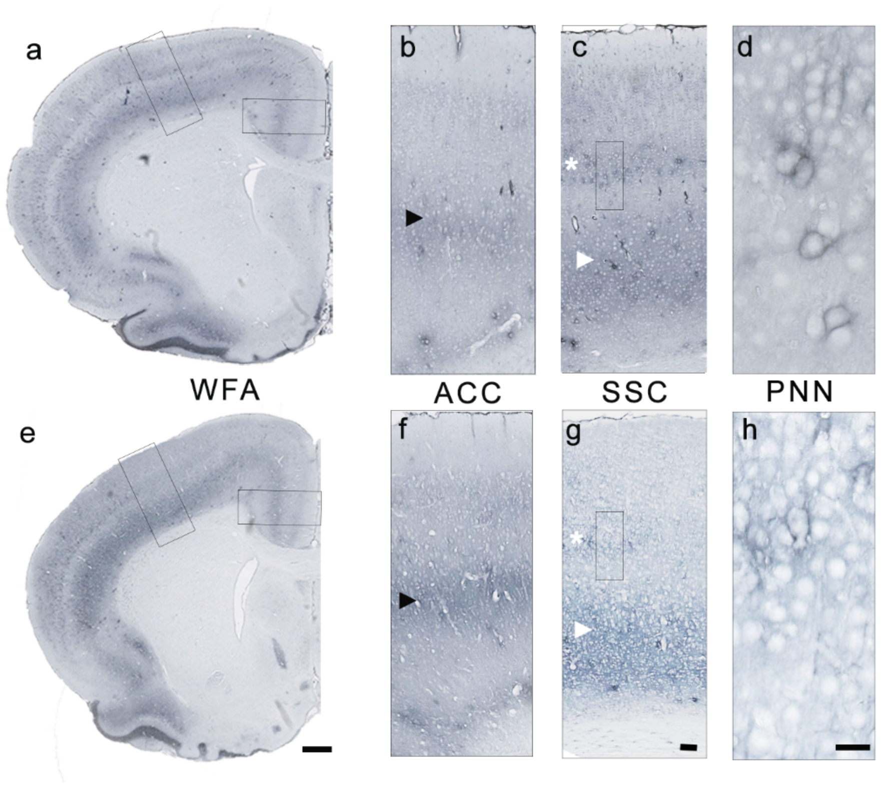


**Supplementary Fig. S6 Lectin-specific glycosylation patterns in control and hypoxic rats at P15**

Coronal sections at the level of the ACC and primary somatosensory cortex (SSC) (~+2.28 mm from Bregma) from control (a–d) and hypoxic rats (e-h), stained with *Wisteria floribunda* agglutinin (WFA).

WFA staining shows preserved glycosylation patterns in the ACC, with the strongest diffuse ECM labeling in layer V (arrowhead in b and f). Black boxes in (a, e, c, g) indicate regions enlarged in (b-h). In the dorsolateral cortex of hypoxic rats, increased diffuse staining is observed in the infragranular layers (arrowhead in g) compared with controls (arrowhead in c). Concurrently, reduced glycosylation is observed in supragranular layers, with less distinct laminar organization in hypoxic rats (asterisk in g). Immature perineuronal nets were observed in both groups (d,h).

Scale bars: 0.5 mm in (e) applies to (a); 50 µm in (g) applies to (b-c; f-g); 25 µm in (h) applies to (d)
