## Supplementary material for "Mild perinatal hypoxia uncouples excitatory–inhibitory circuit maturation and reprograms neocortical organization": Supplementary_tables_bioRxiv.pdf

| EXPERIMENT | AGE | TOTAL NUMBER OF RATS | HYPOXIC | CONTROL | EXCLUDED |
|---|---|---|---|---|---|
| ABS | P1 | 12 (6F; 6M) | 6 (3F; 3M) | 6 (3F; 3M) | |
| BEHAVIORAL TESTING | P3-P14 | 24(12F; 12M) | 12 (6F; 6M) | 12 (6F; 6M) | 3 control males (died due to an unexplained factors during the testing period) |
| IN VIVO MRI | P15 | 16 (16M) | 8 | 8 | DTI (1 control excluded from analysis due to artefacts) |
| IMMUNOHISTOCHEMICAL EXPERIMENTS AND ANALYSIS | P15 | 44 (22F; 22M) | 22 (11F; 11 M) | 22 (11F; 11M) | |
| TOTAL NUMBER | | 56 (28F; 28M) | 28 (14F; 14M) | 28 (14F; 14M) | |

**Supplementary Table 1** Total number of Wistar (RccHan: WIST) rats used in the study

| | T1 map | T2 map | T1-IR | T2w | DTI |
|---|---|---|---|---|---|
| Sequence type | RAREVTR | MSME | RARE | RARE | DTI-EPI |
| Scan dimension | 2D | 2D | 3D | 3D | 3D |
| TR | 550 / 800 / 1200 / 1800 / 2800 / 4000 ms | 3500 ms | 3000 ms | 3000 ms | 1600 ms |
| TE | 12 ms | 12 / 12 / 144 ms | 12 ms | 160 ms | 28 ms |
| Averages | 2 | 2 | 1 | 1 | 1 |
| Sequence-specific parameters | RARE: 2<br>Echo spacing: 6 ms | | RARE: 2<br>Echo spacing: 6 ms<br>Inversion time: 650 ms | RARE: 16<br>Echo spacing: 20 | Diff. directions: 20<br>A0 images: 5<br>B-values: 600 / 1200 s/mm$^2$<br>Preparation: SpinEcho, Double Sampling |
| Acceleration | / | / | / | / | Partial-FT: 1.4 (phase) |
| Bandwidth | 52000 Hz | 15000 Hz | 40000 Hz | 40000 Hz | 348000 Hz |
| Şcan time | 17 min 50 s | 11 min 12 s | 21 min 36 s | 21 min 27 s | 38 min 24 s |

**Supplementary Table 2** Detailed scan parameters

| Primary antibodies | Cat.No | Host, isotype, format | Dilution | Supplier |
|---|---|---|---|---|
| Anti-Microtubule-associated protein-2 (MAP2) | M4403 | Mouse | 1:2000 | Sigma |
| Anti-Neurofilament H (NF-H) Nonphosphorylated; SMI-32 | 801701 | Mouse monoclonal purified, IgG1 | 1:2000 | Biolegend |
| Anti-Parvalbumin PV | ab 11427 | Rabbit polyclonal | 1:2000 | Abcam |
| Anti-Myelin Basic Protein (MBP) | 808401 | Mouse monoclonal | 1:2000 | BioLegend |
| Anti- Complexin 3 | 122302 | Rabbit polyclonal | 1:2000 | Synaptic Systems |
| Anti- Keratan Sulfate (KS), clone 5-D-4 | MABN2483 | Rabbit | 1:150 | MEMD Milipore Corp |
| Lectin from Wisteria floribunda WFA | L1516-2MG | Mouse | 6 ul/ml | Sigma-Aldrich |
| Anti- Lumican | ab 168348 | Rabbit monoclonal | 1:000 | Abcam |
| Anti-Neurocan | N0913 | Mouse monoclonal | 1:600 | Sigma-Aldrich |
| Anti- Versican | PA1-1748A | Rabbit monoclonal | 1:2000 | Thermofisher |

**Supplementary Table 3** List of immunohistochemical antibodies used in the study

| Subject ID | group | mean FA value |
|---|---|---|
| **R6** | 1 | 0.235854611 |
| **R8** | 1 | 0.251721829 |
| **R10** | 1 | 0.240254849 |
| **R12** | 1 | 0.240745664 |
| **R14** | 1 | 0.246833533 |
| **R16** | 1 | 0.235057935 |
| **R18** | 1 | 0.230423659 |
| **R3** | 2 | 0.283537596 |
| **R5** | 2 | 0.266342342 |
| **R9** | 2 | 0.279600680 |
| **R11** | 2 | 0.240789175 |
| **R13** | 2 | 0.237760618 |
| **R15** | 2 | 0.284678876 |
| **R17** | 2 | 0.266762614 |
| **R19** | 2 | 0.257465929 |
| **MEAN± SD GROUP 1 (CONTROLS)** | 0.2401 ± | 0.0073 |
| **MEAN± SD GROUP 2 (HYPOXIC)** | 0.2646 ± | 0.0183 |
| **P-VALUE (t-test)** | 0.0064 | |

**Supplementary Table 4** mean FA values in ROI (ACC-significant cluster)
